# Dynamics of duplicated gene regulatory networks governing cotton fiber development following polyploidy

**DOI:** 10.1101/2024.08.12.607624

**Authors:** Xianpeng Xiong, De Zhu, Corrinne E. Grover, Jonathan F. Wendel, Xiongfeng Ma, Guanjing Hu

## Abstract

Cotton fiber development entails complex genome-wide gene regulatory networks (GRN) that remain mostly unexplored. Here we present integrative analyses of fiber GRNs using public RNA-seq datasets, integrated with multi-omics genomic, transcriptomic, and cistromic data. We detail the fiber co-expression dynamics and regulatory connections, validating findings with external datasets and transcription factor (TF) binding site data. We elucidate previously uncharacterized TFs that regulate genes involved in fiber-related functions and cellulose synthesis, and identify the regulatory role of two homoeologous G2-like transcription factors on fiber length. Analysis of duplicated gene expression and network relationships in allopolyploid cotton, which has two co-resident genomes (A, D), revealed novel aspects of asymmetric subgenomic developmental contributions. Whereas D-based homoeolog pairs drive higher overall gene expression from the D subgenome, TFs from the A subgenome play a preferential regulatory role in the fiber gene regulatory network. Following allopolyploid formation, it appears that the trans-regulatory roles of TFs diversified more rapidly between homoeologs than did the cis-regulatory elements of their target genes. Our approach underscores the utility of network analysis for detection of master regulators and provides fresh perspectives on fiber development and polyploid functional genomics, through the lens of co-expression and GRN dynamics.

## Introduction

Cotton ranks among the world’s most important agricultural plants, supplying most of our natural textile fibers. The remarkable cotton “fibers”, which are extensively elongated and naturally twisting single cells originating from the ovule epidermis, undergo a complex developmental program, entailing five sequential yet overlapping stages: initiation, elongation, transition, secondary cell wall (SCW) thickening, and maturation (Haigler et al. 2012). These complex, coordinated stages are crucial for fiber production, as initiation determines the number of epidermal cells that develop into fibers, while elongation and SCW thickening determine the final length and strength of each fiber (Yang et al. 2014). Given the importance of these stages, the past two decades have witnessed considerable progress towards elucidating the principal pathways and genes orchestrating fiber initiation and elongation, mostly focusing on the regulatory role of transcription factors (TF) (Huang et al. 2021).

The initiation of cotton fibers shares a similar mechanism with the development of *Arabidopsis* leaf trichome, regulated by the intricate MYB-bHLH-WDR (MBW) transcriptional complex (Wang et al. 2019; Zhang et al. 2019; Wen et al. 2023). A specific MIXTA-like MYB TF, *GhMYB25-like*, serves as a pivotal switch in this context, with suppression leading to abnormal fiber cell initiation and fiberless seeds (Walford et al. 2011). The elongation phase of cotton fiber development is distinctive among plant cells, involving special factors and mechanisms that confer extraordinary fiber length and growth rate. Many TFs, including HD-ZIP, TCP, WRKY, and ARF, are integral to modulating this phase (Wen et al. 2022). Notably, the cotton HD-ZIP family TF *GhHOX3* promotes fiber elongation by upregulating transcription of the cell wall loosening protein genes *GhRDL1* and *GhEXPA1* (Shan et al. 2014), while a fiber-preferential WRKY TF GhWRKY16 directly activates the transcription of *GhHOX3*, a MYB family TF (*GhMYB109*), and a cellulose synthase gene (*GhCesA6D-D11*) (Wang et al. 2021b). TEOSINTE BRANCHED, CYCLOIDEA AND PCF 14 (GhTCP14) mediates cotton fiber elongation by directly activating the expression of auxin-responsive gene *GhIAA3* and auxin transporter genes *GhPIN2* and *GhAUX1* (Wang et al. 2013). Beyond their involvement in elongation, *GhHOX3, GhTCP14*, and *GhWRKY16* have also been confirmed to positively regulate fiber initiation (Qin et al. 2022; Wen et al. 2023). Transitioning from elongation to SCW thickening, several TFs, such as *GhFSN1* (Zhang et al. 2018) and five MYB family TF (*GhMYB1* (Yadav et al. 2017), *GhMYBL1* (Sun et al. 2015), *GhMYB7* (Huang et al. 2016), *GhMYB46_D9*, and *GhMYB46_D13* (Huang et al. 2019) have been reported to positively regulate SCW thickening. *GhTCP4* and a Class II KNOX TF (*GhKNL1*) function both in fiber elongation and SCW thickening; however, interestingly, GhKNL1 represses genes promoting elongation and SCW cellulose deposition, whereas GhTCP4 coordinates the suppression of fiber elongation through its interaction with *GhHOX3* to activate SCW synthesis (Cao et al. 2020; Wang et al. 2022). These results underscore the nuanced regulatory roles of TFs and intricate dynamics across different stages. Despite extensive research into TF-mediated regulatory networks of cotton fiber development, these studies have been primarily conducted in a gene-by-gene fashion, leaving relatively unexplored a more comprehensive understanding of the dynamic interactions among networks of genes governing fiber development.

Organ development in plants is intricate, depending on the precise timing and spatial regulation of gene expression, a process captured by complex gene regulatory networks (GRNs) (Haque et al. 2019; Jones and Vandepoele 2020; Vandepoele and Kaufmann 2023). These networks represent the full suite of interactions between TFs and their target genes, where TFs bind to specific DNA sequences known as TF binding sites (TFBSs) and regulate the transcription of downstream targets. Central to GRNs are hub TFs, which, due to their large number of target and/or regulating genes, are crucial for the integration and dissemination of regulatory signals across the network (Barabási and Oltvai 2004; Levine and Davidson 2005). Identifying hub genes in plant GRNs offers a clear roadmap for pinpointing master regulators and unraveling interconnections essential for biological processes and developmental programs (Gaudinier and Brady 2016; Haque et al. 2019; Jones and Vandepoele 2020). These network components, when modulated, can enhance plant productivity or resilience, often yielding more significant influence over complex phenotypes than manipulating individual genes alone (Springer et al. 2019). Therefore, the construction and mining of GRNs is key for increasing the predictive power of genome engineering approaches aimed at agronomic traits for crop improvement.

GRN construction methods can be broadly categorized into two main approaches differentiated by the source of information utilized: data-driven methods and prior knowledge-based methods. Data-driven methods leverage high-throughput experimental techniques to unveil physical interactions between TFs and their target genes. These techniques includes: (1) Chromatin immunoprecipitation sequencing (ChIP-seq) (Furey 2012) which identifies genomic sites bound by a given TF *in vivo*; (2) DNA-affinity purification sequencing (DAP-seq) (O’Malley et al. 2016) captures DNA bound by the *in vitro* expressed TF; and (3) yeast one-hybrid assay (Taylor-Teeples et al. 2015), which identifies physical interactions between TFs and their potential DNA binding sites. Additionally, chromatin accessibility assays (Song and Crawford 2010; Buenrostro et al. 2015; Zhao et al. 2020), including DNase-I hypersensitive site sequencing (DNase-seq), assay for transposase-accessible chromatin with sequencing (ATAC-seq), and MNase hypersensitive sequencing (MH-seq), have also been applied to characterize *cis*-regulatory elements as potential transcription factor binding sites (TFBSs) at a genome-wide scale, thereby revealing regulatory relationships between TFs and target genes. Despite the substantial increase in information about regulatory sequences and interactions offered by these assays, inherent technical challenges and cost still pose limitations for studying large numbers of TFs (Kulkarni and Vandepoele 2020). Consequently, only a few plant species, such as *Arabidopsis* and maize, have constructed GRNs based on large-scale experimental data of regulatory interactions (Taylor-Teeples et al. 2015; Gaudinier et al. 2018; Tu et al. 2020; Tang et al. 2021).

In contrast to data-driven methods, prior knowledge-based methods for GRN construction integrate existing biological knowledge, drawing from scientific literature, known biological pathways, functional gene ontology categories, and various knowledge databases of gene-to-gene relationships (Linde et al. 2015). For instance, resources like PlantTFDB (https://planttfdb.gao-lab.org/) and PlantRegMap (https://plantregmap.gao-lab.org/) serve as integrated platforms for plant regulatory data and analysis, which systematically screens for functional TFBSs and regulatory interactions in plants. PlantRegMap, in particular, curates additional functional and evolutionary annotations, such as expression profiles and multiple-species comparisons, along with corresponding literature references, resulting in generation of regulatory maps for the main lineages of angiosperms, effectively representing their preliminary GRNs (Tian et al. 2020a). However, these general GRNs do not account for differences in gene regulation relationships across different tissues, developmental stages, or conditions.

Integration methods often combine both data-driven and prior knowledge-based approaches, leveraging expression data underlying specific states to refine GRNs built based on existing knowledge, or vice versa. Computational algorithms that infer GRNs from gene expression data include correlation and information theory-based methods, probabilistic graphical models, and machine learning (Haque et al. 2019). Correlations and mutual information methods assume that co-expression is an indicator of coregulation and deterministically controlled by upstream regulators. Probabilistic graphical models consider gene expression as random variables with a certain probability distribution over different tissues and conditions. Machine learning algorithms, such as ensemble decision trees and support vector machines, are trained on expression data to predict regulatory relationships between genes. In recent years, these inference methods have been employed to construct GRNs and identify important genes and regulatory relationship involved in plant growth and developmental processes, such as photomorphogenesis in *Arabidopsis* (Balcerowicz et al. 2021), abiotic and disease responses in wheat (Ramírez-González et al. 2018), nitrogen-deficiency responses in rice (Ueda et al. 2020), as well as Kranz anatomy development in maize and rice (Chang et al. 2019). More recently, as demonstrated for spike phenotypic traits in wheat and flowering time regulation in maize (Chen et al. 2023; Han et al. 2023), GRN inference has been improved by integrating heterogeneous -omics and functional validation data for a more comprehensive understanding of the biomolecular networks. Despite the critical importance of cotton fiber development to its success as a major crop species, a comprehensive GRN that unravels the intricate molecular mechanisms underlying fiber traits is still lacking.

In this study, we employed three distinct inference methods to construct GRNs utilizing transcriptome data from 401 samples. Notably, we validated the robustness and efficacy of resulting GRNs through rigorous integration with prior knowledge-based regulatory maps, DAP-seq data, and additional transcriptomic datasets from gene perturbation experiments. Through this integrative analysis, we identified novel transcription factors crucial for orchestrating fiber development. We further validated the functional significance of a homoeologous pair of top-ranked G2-like TF genes (*GhMYS1_A10* and *GhMYS1_D10*) in the GRN, revealing their potential regulatory mechanisms in fiber development.

An additional important dimension of our study is that it addresses the fate of duplicated GRNs in an allopolyploid plant, that is, one that contains two co-resident genomes. *Gossypium hirsutum* contains the descendant genomes of both its A-genome and D-genome ancestors (each n=13), and thus has an AD-genome with an additive (n=26) chromosome number. This evolutionary history raises the possibilities of revealing the fate of duplicated GRN dynamics following allopolyploid evolution, a prominent process in plant evolution (Hu et al. 2021; Viot and Wendel 2023). Here, we elucidate subgenomic control over fiber expression at both the genic co-expression and GRN levels, providing insights into the regulatory landscape of fiber development in an allopolyploid contect. Finally, we provide an integrative network resource and demonstrate its utility in enhancing our understanding of cotton fiber development, thereby facilitating targeted interventions to modulate fiber traits.

## Results

### A fiber gene expression atlas of the Upland cotton *G. hirsutum*

We compiled a dataset of 473 Upland cotton (*G. hirsutum*) fiber transcriptomes from 12 RNA-seq studies (Tuttle et al. 2015; Zhang et al. 2015, 2021a; Hinchliffe et al. 2016; Lu et al. 2017; Bao et al. 2019; Hu et al. 2019; Sun et al. 2019a; Huang et al. 2020; Li et al. 2020; He et al. 2021) (Supplementary Table S1). These samples spanned key fiber developmental stages from 0 to 30 days post-anthesis (dpa), including fiber initiation, elongation, transition, and secondary cell wall (SCW) synthesis (Figure 1A, Supplementary Fig. S1). To ensure specificity to fiber cells, 12 samples from 0 to 3 dpa obtained from whole ovules were excluded. After quality screening based on a unique mapping rate higher than 70% and outlier removal through principal component analysis (PCA), a final set of 413 high-quality samples was obtained with Q20 above 93.09% (Supplementary Table S2). Further refinement using principal component analysis (PCA) and t-distributed stochastic neighbor embedding (t-SNE) led to the removal of another 12 outlier samples, resulting in a final dataset of 401 samples (Supplementary Fig. S2, Supplementary Table S2).

**Figure 1.**
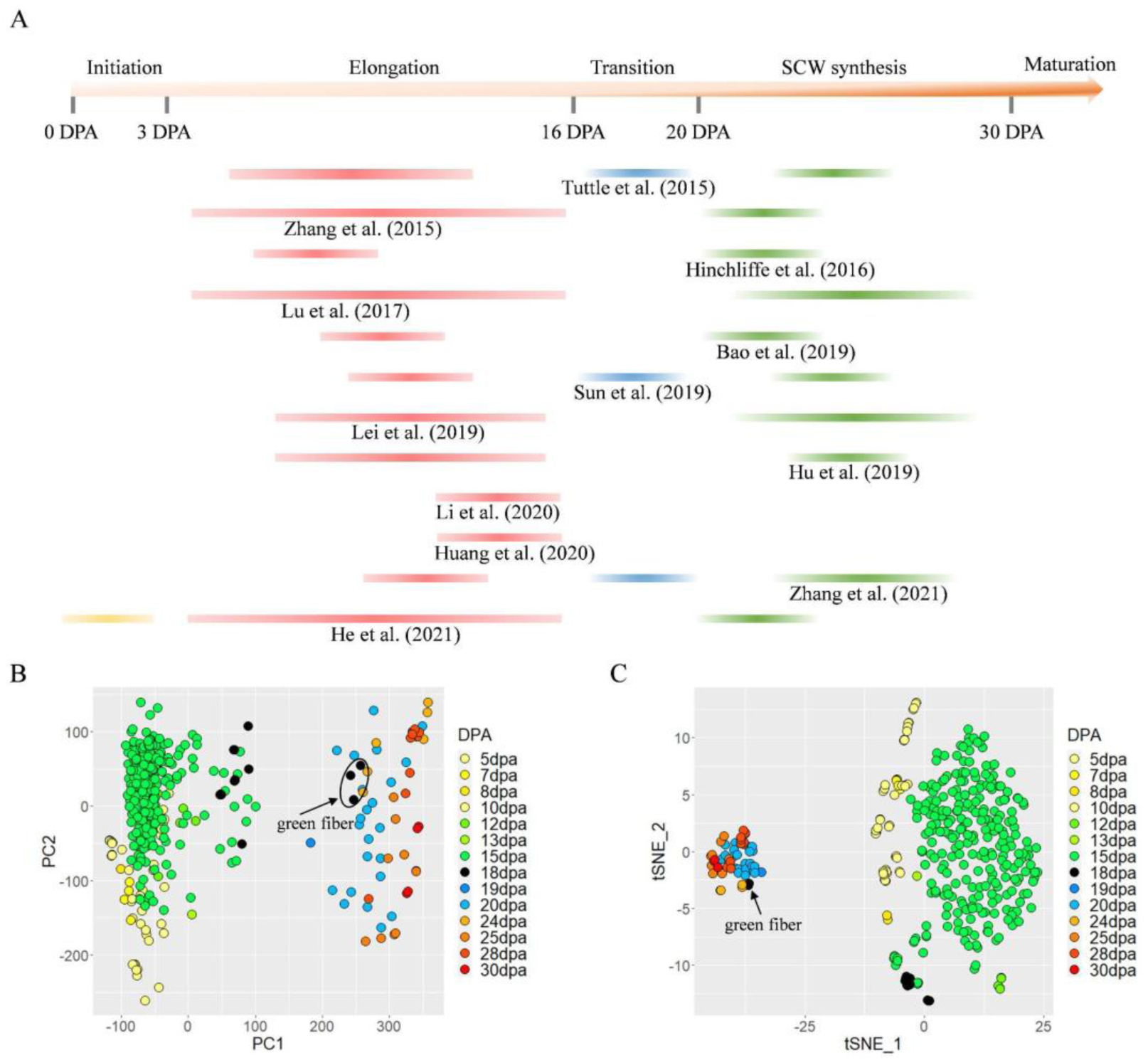
Cotton fiber transcriptomic datasets for this study. (**A**) Timeline displaying the stages represented by the 12 studies used to generate a dataset of 401 fiber RNA-seq samples for an in-depth exploration of cotton fiber development. Fiber elongation, transition, and SCW synthesis stages are indicated by red, blue, and green bars, respectively, and each line represents one existing dataset. This color scheme is applied consistently across all figures here. (**B**) Principal component analysis (PCA) of 57,151 gene expression profiles. PC1 and PC2 captured 16.8% and 11.5% of variance, respectively. (**C**) T-distributed stochastic neighbor embedding (t-SNE) was also employed for dimension reduction and visualization of the fiber expression landscape.

Based on the standardized gene expression by TPM (transcripts per million), both PCA t-SNE identified two distinct clusters of fiber samples: one comprising 329 samples from 5 to 15 dpa and another with 57 samples from 19 to 30 dpa, while 15 samples from 18 dpa exhibited an intermediate distribution (Figure 1B, C). This observation indicates the pronounced transcriptional distinction of the fiber cell from about 19 dpa as it becomes intensely committed to SCW synthesis. Categorization of the three earlier stages was less clear, likely due to genetic variation and variation in growing conditions or collection techniques across studies. Notably, the inclusion of natural green-fiber cotton varieties highlights developmental differences that can distinguish accessions. That is, among the 15 samples representing 18 dpa fiber, the 12 samples derived from white-fiber producing accessions clustered with the 5 to 15 dpa samples (circa 50 on PC1), whereas the 3 green-fiber (variety Xincai 7) samples clustered with the 19 to 30 dpa samples, suggesting that the green-fiber accessions transition to SCW synthesis sooner than the white-fiber accessions represented here and underscoring the potential for temporal differences in development among cotton varieties.

To ensure the reliability of this dataset, we examined expression patterns of 192 fiber-related genes known for their roles in cotton and/or *Arabidopsis* trichome development (Supplementary Table S3). These genes were classified into three groups based on their average TPM values per dpa (Supplementary Fig. S3). Group I, comprising 93 genes, displayed high expression levels early during fiber elongation, featuring well-known elongation-associated genes like *GhMYB25* (Machado et al. 2009), *GhMYB25-like* (Walford et al. 2011), *GhTCP4* (Cao et al. 2020), *GhPIN3a* (Zeng et al. 2019), *GhHOX3* (Shan et al. 2014), *GhHD1* (Walford et al. 2012), *GhCaM7* (Tang et al. 2014), *GhWRKY16* (Wang et al. 2021b), and *GhBZR1* (Zhou et al. 2015). Group II, containing 29 genes exhibiting higher expression during SCW synthesis at later time points, included established SCW genes such as *GhBZR3* (Shi et al. 2022), *GhKNL1* (Gong et al. 2014), *GhSWEET12* (Sun et al. 2019b), *GhFSN1* (Zhang et al. 2018), and *GhMYB46_D13* (Huang et al. 2019). Group III comprised 70 genes with expression profiles peaking at various time points between 5 and 30 DPA. The expression patterns observed here closely align with previous reports (Supplementary Table S3); that is, 77% (105 out of 136) of the genes surveyed exhibited the expected expression profiles, providing robust validation of our gene expression atlas. The few inconsistencies observed were primarily attributed to missing data (i.e., lack of later time point data or large gaps between time points) in earlier studies. For example, several genes (including *GhGA20ox1* (Xiao et al. 2010), *GhTUA9* (Li et al. 2007), *GhMYB212* (Sun et al. 2019b), *GhACO1* (Wei et al. 2022), *GhMAH1* (Ma et al. 2022), *GhMYB5_A12* (Wang et al. 2021a), and *GhCPC* (Liu et al. 2015)), which were previously compared only between 5 and 15 dpa, exhibited continuous expression changes in 5 to 30 dpa based on our comprehensive expression profiles. Additionally, our dataset revealed that several well-known fiber initiation genes, including *GhMYB25* (Machado et al. 2009), *GhPIN6* (Zhang et al. 2017b), *GhPIN3a* (Zeng et al. 2019), *GhBZR3* (Shi et al. 2022), and *GhSWEET12* (Sun et al. 2019b), exhibited high expression levels in later stages of development that were not previously examined. This suggests that these genes may have regulatory roles beyond fiber initiation, highlighting insights enabled by our comprehensive data analysis.

### Co-expression gene network analysis reveals fiber developmental dynamics

To explore the transcriptional dynamics of cotton fiber development, we employed weighted co-expression gene network analysis (WGCNA) on fiber-expressed genes. Opting for a filtering criteria of TPM>0 in 30% of the samples (see Methods and Supplementary Fig. S4), we identified and included 57,151 genes for further analysis, representing 76.3% of the total genome expressed in fibers, consistent with previous reports (Hovav et al. 2008a; Yoo and Wendel 2014; Gallagher et al. 2020). The subsequent WGCNA analysis categorized 34,075 genes into 20 co-expression modules, varying in size from 109 to 7,360 module gene members (Supplementary Fig. S5). The seven largest modules, ME1 (turquoise), ME2 (blue), ME3 (brown), ME4 (tan), ME5 (green), ME6 (black), and ME7 (red), collectively accounted for 87.7% (29,884 genes) of all co-expressed genes. The remaining 23,076 genes, which could not be assigned to any modules, were grouped into a grey module, indicating no discernible co-expression relationships.

To examine phenotypic associations, we correlated module eigengenes (MEs) with fiber development for 14 time points between 5 and 30 dpa (inclusive; Figure 2A). Pearson correlation analysis showed significant associations with fiber development for the majority of modules, treating the time points as a binary categorical variable (Figure 2A) or a numeric variable (DPA; Supplementary Fig. S5B). ANOVA of MEs revealed significant developmental changes for seventeen modules, excluding ME6, ME17, and ME20 (MEs ∼ DPA, ANOVA *P* < 0.05) (Figure 2B).

**Figure 2.**
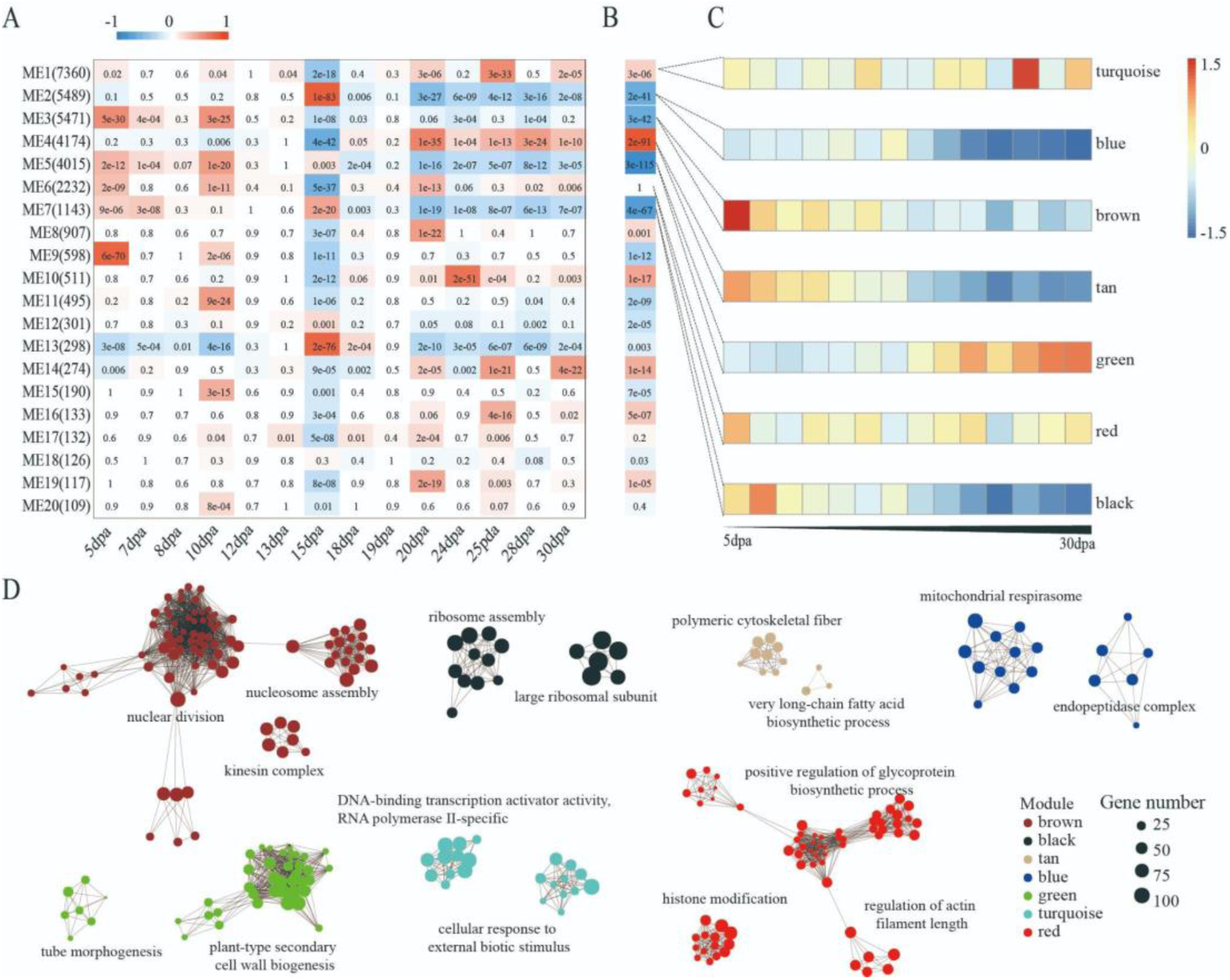
Phenotypic and functional associations of co-expression gene modules during fiber development. **(A)** For the 20 co-expression gene modules identified by weighted gene co-expression network analysis (WGCNA), heatmap represents Pearson correlation coefficients and *P*-values (cell color and text, respectively) between the module eigengenes (MEs, by row) and fiber developmental stages treated as the binary categorical variable (by column). (**B**) ANOVA of MEs (by column) by fiber developmental stages treated as a numeric variable (MEs, by row). Heatmap cell color and text represent Pearson correlation coefficients and *P*-values, respectively. **(C)** Heatmap of z-score normalized MEs for the seven largest modules across fourteen fiber developmental time points. (**D**) Gene Ontology (GO) enrichment analysis of the seven largest modules, displaying the top two most significant interconnected GO clusters terms each. Different colors represent corresponding modules.

Notably, the green module displayed the highest correlations with DPA (*r* = 0.80, *P* = 2e-91), exhibiting a gradually increasing expression profile along fiber development. This module consists of 4015 genes and 223 TFs, and it was enriched with GO terms related to cell wall development, such as plant-type secondary cell wall biogenesis, cell wall polysaccharide biosynthetic process, hemicellulose metabolic process, tube morphogenesis, and cell wall macromolecule biosynthetic process (Figure 2D). Conversely, the brown, blue, and tan modules showed strong negative correlations (*r* = −0.61∼-0.85, *P* < 2e-41), corresponding to decreasing expression along fiber development (Figure 2B-C). The tan module in particular showed significant enrichment of GO terms associated with cotton fiber development, encompassing processes like very long-chain fatty acid metabolism, microtubule organization, cell tip growth, pectin biosynthesis, and polymeric cytoskeletal fiber processes (Figure 2D; Supplementary Fig. S6; Supplementary Table S4).

The turquoise module, the largest with 7360 genes (including 984 TFs), peaked in expression at 25 dpa. Significant enrichment with GO terms including core promoter sequence-specific DNA binding, DNA-binding transcription activator activity, and RNA polymerase II-specific were observed. The red module, consisting of 2232 genes (including 50 TFs) and without significant correlation with DPA, were enriched with diverse GO functions (Supplementary Table S4).

Exploring the roles of hormone signaling pathways in regulating cotton fiber development (Huang et al. 2021), we observed significant functional enrichment in key modules. The brown module with expression peaking at 5 dpa revealed a strong association with auxin (IAA)-activated signaling pathways (Supplementary Fig. S7A), consistent with the known function of IAA-activated signaling pathways in promoting fiber initiation and elongation. Surprisingly, brassinosteroid (BR)-related signaling pathways, known to regulate fiber initiation and elongation, were enriched in both the tan module (early peaking at 5 dpa) and the turquoise module (late peaking at 25 dpa). This introduces a novel perspective on the impact of BR post-fiber elongation, which has not been reported previously (Supplementary Fig. S7B-C). Additionally, the red and turquoise modules, which exhibited more complex and dynamic gene expression patterns across development, were enriched for BR, jasmonic acid, gibberellin, and ethylene-related signaling pathways (Supplementary Fig. S7C), warranting further investigation into their functional implications. Besides these extensively studied phytohormones, the turquoise module also showed significant enrichment of abscisic acid, cytokinins, and salicylic acid-related signaling pathways (Supplementary Fig. S7D), While these pathways are recognized for their roles in plant growth and development (Santner and Estelle 2009), their specific impact on cotton development remains underexplored.

### Construction and evaluation of the cotton fiber gene regulatory networks

To infer regulatory interactions beyond co-expression relationships between genes, we systematically constructed gene regulatory networks (GRNs) using three distinct inference methods: Corto, GENIE3, and dynGENIE3. Leveraging the 57,151 fiber-expressed genes derived from 401 RNA-seq samples, we evaluated the regulatory relationships between 3,638 transcription factors (TFs) and their putative target genes. Both GENIE3 and dynGENIE3 inferences were confined to the top one million TF-target interactions (edges) for comparison, retaining over twice the number of genes (nodes) from the GENIE3 network than from the dynGENIE3 network (54,237 and 25,441, respectively). Although Corto inferred only 232,943 TF-target interactions (edges), it retained a comparable number of nodes to GENIE3 (56,052), resulting in the densest and most clustered network topology among the three methods, followed by GENIE3 and then dynGENIE3 (Table 1). Because differences in GRN construction can lead to different inferences, we evaluated GRN quality for each method based on existing and newly generated data, as listed below.

**Table 1.**
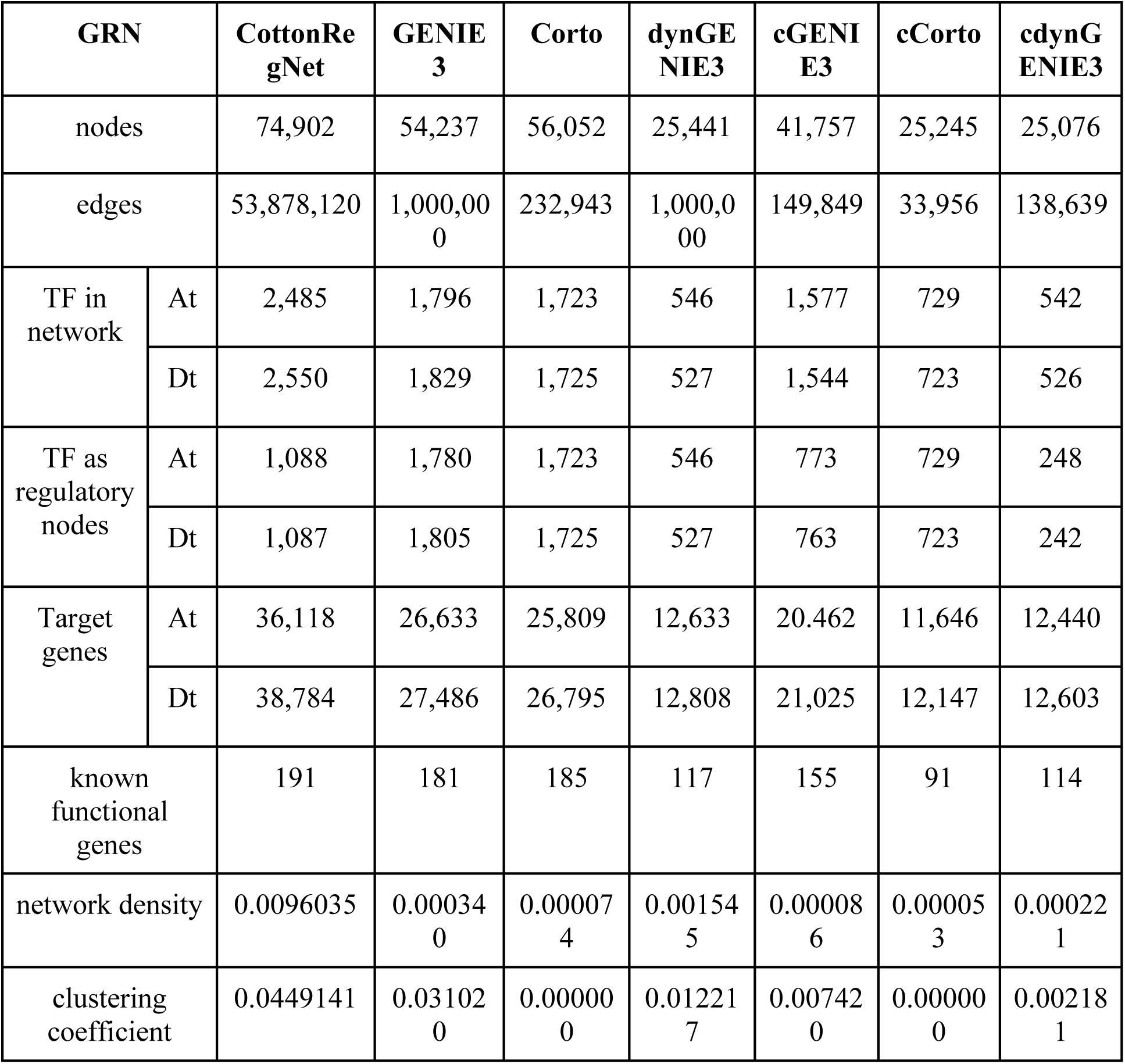
Fiber gene regulatory networks constructed.

We first assessed the ability of each GRN to capture documented TF-target interactions based on systematic literature mining in plants, as assembled into the PlantRegMap (Jin et al. 2015). These known regulatory relationships were projected onto cotton orthologs to generate the cottonRegMap. Among the three GRN methods, GENIE3 outperformed Corto and dynGENIE3, recovering the highest percentage of known interactions (14.98% vs. 14.58% and 13.85%, respectively), although the range among these percentages is relatively small. We also note that these seemingly low percentages of interactions reflect the non-specific nature of cottonRegMap, which involves prior knowledge assembled from various plants and is not specific to cotton fibers. Without a true gold-standard dataset for validation, we employed a permutation test to determine the expected number of interactions captured by chance. Both GENIE3 and Corto captured more interactions than the expected 14.37% of interactions (bootstrapping *P* < 0.05), demonstrating their utility in capturing biological information for cotton fiber (Figure 3A). For subsequent analyses, we integrated prior biological knowledge by retaining only the regulatory interactions predicted by GENIE3, dynGENIE3, and Corto that were also supported by cottonRegNet. This approach allowed us to retain the relative topological patterns between methods (Table 1). These integrated networks were designated as cGENIE3, cdynGENIE3, and cCorto, respectively.

**Figure 3.**
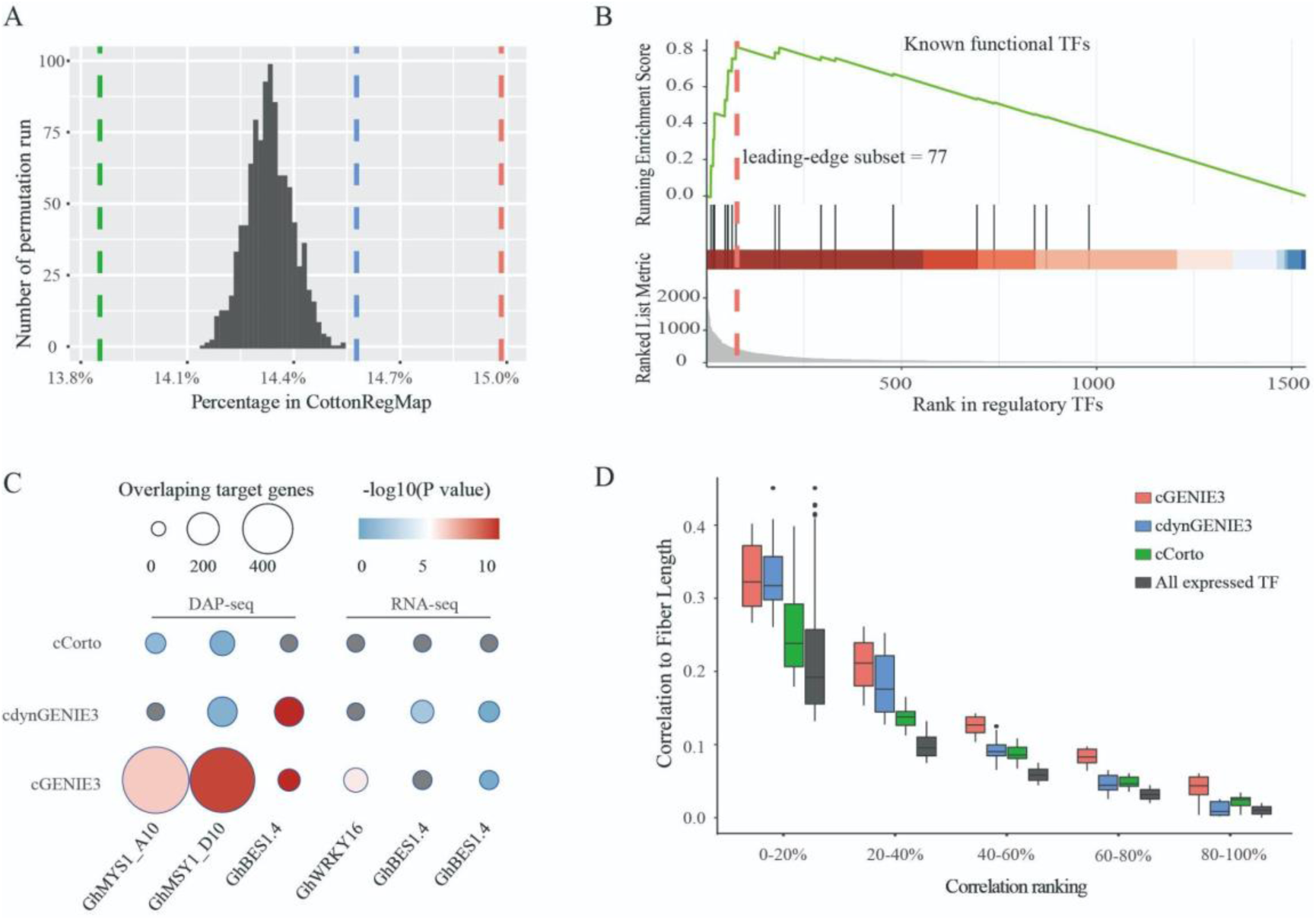
Evaluation of fiber GRN inferences. (**A**) Histogram presents the bootstrap distribution (n=1000) of cottonRegMap TF-target relationships as captured by chance. Red, blue, and green lines represent the cottonRegMap TF-target relationships inferred by GENIE3, dynGENIE3, and Corto, respectively. Both GENIE3 and Corto inferred significantly more interactions outside the bootstrap distribution. (**B**) GSEA of known functional TFs among TFs rankings inferred by cGENEI3. The enrichment score reflects the degree of over-representation of a set of 54 known functional TFs at the top of the ranked TFs identified by cGENIE3. The red dashed line indicates that these known functional TFs were significantly enriched at the top 77 ranking TFs. (**C**) Heatmap of overlapping target genes between empirical evidence (columns) and GRN inferences (rows). WRKY16, with GRN inferences for cGENIE3, cdynGENIE3, and cCorto. Each cell represents the number of overlaps and the significance of the corresponding hypergeometric test. DAP-seq results of *GhMYS1_A10*, *GhMYS1_D10*, and *GhDES1.4* and RNA-seq results of *GhDES1.4* and *GhWRKY16* were shown. (D) The correlation between expression variation of 77 hub TFs and fiber length was significantly higher than that of 3,638 TFs expressed in fibers. Five different percentages ranks were divided according to the correlation between TF and fiber length, where 0% to 100% represent increasing correlation.

Using a second approach, we assessed the ability of each GRN to recover known fiber-related functional genes and TFs (Supplementary Table S3) previously reported in the literature. We curated 192 fiber-related functional genes, of which the cGENIE3 network contained 155 (80%), the cdynGENIE3 network contained 114 (59%), and the cCorto network contained 91 (47%) of the genes on the list. In terms of the percentage of known genes among total network nodes, cdynGENIE3 exhibited the highest percentage (0.45%, 117 of 25,441), followed by cGENIE3 (0.37%, 155 of 41,757) and cCorto (0.36%, 91 of 25,245). We extended this to assess whether known TFs were enriched among the highly ranked TF regulators in each GRN. Gene set enrichment analysis (GSEA) showed that the curated TFs were significantly enriched at the top of the cGENIE3 network; specifically, a leading-edge subset comprising 77 TFs was identified as the most significant contributors to this enrichment (Figure 3B, Supplementary Table S5). In contrast, known TFs were not enriched at the top of cdynGENIE3 and were randomly distributed in rank in the cCorto network (Supplementary Fig. S8). These results suggested that cGENIE3 has stronger prediction power for key TFs compared to cdynGENIE3 and cCorto.

We further validated the GRN-inferred TF-target relationships for two top ranked (homoeologous) TFs using physical evidence from DNA-affinity purification sequencing (DAP-seq), an *in vitro* genome-wide assay of TF-DNA binding (O’Malley et al. 2016). The homoeologous G2-like TFs *GhMYS1_A10* (*Gohir.A10G036400*) and *GhMYS1_D10* (*Gohir.D10G037100*) were among the most confident (highest-ranked) regulators in all three networks; therefore, these genes, were independently assayed for genome-wide binding sites using DAP-seq (Figure 3C, Supplementary Table S6). These assays yielded 227,117 and 141,945 peaks for *GhMYS1_A10* and *GhMYS1_D10*, respectively, with approximately 8.27% and 6.73% of the peaks located within 2kb of the transcription start site for 10,132 and 9,363 genes (Supplementary Fig. S9A-D). Among these genes, 7,784 and 6,773 were expressed in fibers and identified as targets for *GhMYS1_A10* and *GhMYS1_D10*, respectively (Supplementary Fig. S9F-E). Examination of the overlap in target genes between DAP-seq and each GRN revealed a significant association for both TFs in cGENIE3 (hypergeometric test *p*-values of 2.24e-08 and 3.56e-06) and for *GhMYS1_D10* only in cdynGENIE3 (*p* = 0.0471); no significant association was found for the cCorto GRN (Figure 3C, Supplementary Fig. S9F-G). These results were reiterated when we compared DAP-seq for an additional gene (*GhBES1.4*) with each GRN. *GhBES1.4* is a known core TF in the BR signaling pathway that positively regulates fiber elongation (Liu et al. 2023), yet it was ranked eighteenth by different GRN methods. Using a published DAP-seq dataset for *GhBES1.4* (Liu et al. 2023), we found significant overlaps between the 1214 fiber-expressed target genes of *GhBES1.4* inferred by DAP-seq (Supplementary Fig. S9G) and the cGENIE3 and cdynGENIE3 GRNs, but not the cCorto GRN (Figure 3C).

Our fourth approach utilized published RNA-seq data from TF mutants or transgenic lines to evaluate the accuracy of GRN inferences, by assessing how well the predicted regulatory interactions in the GRNs corresponded to the differentially expressed genes (DEGs) observed in these TF mutants or transgenic lines. Specifically, we identified 3,508 DEGs in *GhWRKY16* RNAi lines, 1,422 in *GhBES1.4* RNAi lines, and 1,790 in *GhBES1.4* overexpression lines compared to wild-type plants, most of which (96.2-99.0%) were expressed in the fiber dataset evaluated here (Supplementary Fig. S10A-C). These DEGs likely represent downstream targets of the TFs perturbed in each respective experiment, and are thus useful to validate our GRN predictions. For *GhWRKY16*, which is a WRKY TF known for promoting fiber initiation and elongation (Wang et al. 2021b), we found significant overlap between the 3,472 DEGs identified from the RNAi line comparison and the *GhWRKY16*-target relationships found in the cGENIE3 GRN (hypergeometric test *p* = 3.90e-06; Figure 4C); in contrast, cCorto and cdynGENIE3 inferred only two and zero DEG targets, respectively. Conversely, the 1,477 DEGs detected in the *GhBES1.4* exhibited significant overlap with the *GhBES1.4*-targets only recovered for the cdynGENIE3 GRN (*p* = 0.005751457). Notably, no significant overlap was found between the DEGs from the *GhBES1.4* RNAi lines and any of the networks (Figure 3C). Combined with the DAP-seq evaluation, these results suggest that both GENIE3 and dynGENIE3 outperform Corto in predicting regulatory targets for specific TFs, notwithstanding the inherent variance depending on the TF and experimental context.

**Figure 4.**
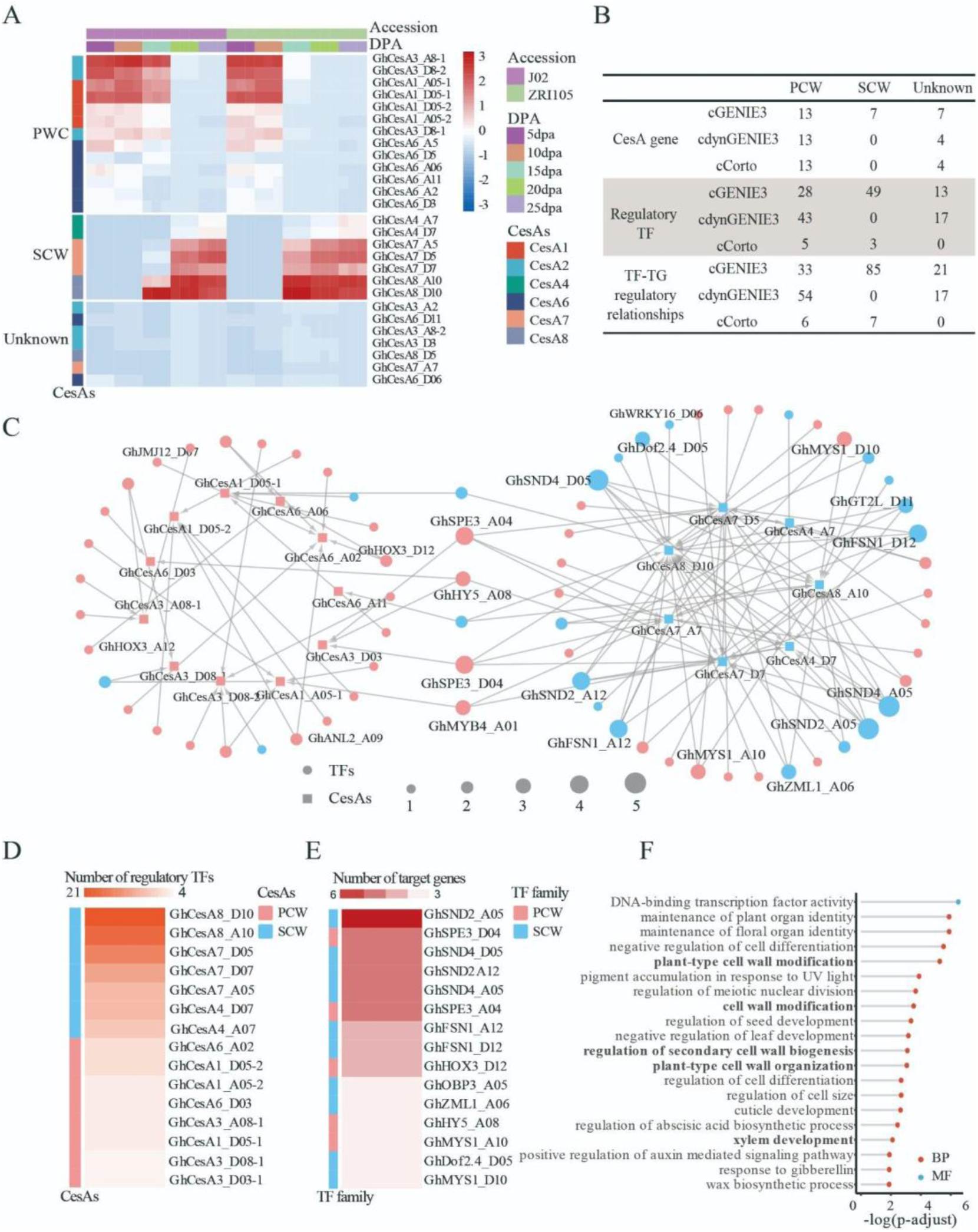
GRN performance in cotton cellulose synthesis. (**A**) Categorization of *GhCesAs* based on gene expression patterns during cotton fiber development. Heatmap presents TPM expression levels in the long-fiber variety J02 and the short-fiber cotton variety ZRI015. Three hierarchical clusters correspond to PCW-related, SCW-related, and unknown *GhCesAs.* (**B**) The number of CesA genes, regulator transcription factors (TFs), and regulatory relationships identified by cGENIE3, cdynGENIE3, and cCorto. (**C**) Cellulose synthesis-related subnetwork inferred by cGENIE3. Square and round nodes represent *GhCesAs* and TFs, respectively, which are connected by directed edges indicating the TF-target relationships inferred. Red and blue node colors represent the categorization of PCW-related and SCW-related genes based on expression patterns during fiber development, respectively. Two network components were detected corresponding to PCW (left) and SCW (right), which were co-regulated by six TFs in the middle. (**D**) Ranking *GhCesAs* by in-degree (i.e., number of incoming linking) from all TFs inferred by cGENIE3. (**E**) Ranking cellulose synthesis related TFs by out-degree (i.e., number of outward links) to target *GhCesAs*. (**F**) Enriched GO terms associated with the 71 TFs inferred by cGENIE3.

Our final assessment correlated the trait fiber length with key fiber TFs inferred by the GRNs. Using the top 77 TFs ranked by each GRN method, Pearson correlation analysis between their expression levels in 15 DPA fiber and mature fiber length revealed the highest phenotypic correlations were found with TFs implicated in the cGENIE3 network, followed by cdynGENIE3 and then cCorto (Figure 3D, Supplementary Table S5, 7-8). All networks showed significantly higher correlations with phenotype than did all 3638 TFs expressed in fibers (Figure 3D).

### Performance evaluation of GRN inferences in the case of cotton cellulose synthesis

In this case study, we evaluated three GRN methods by focusing on their ability to predict regulatory relationships involved in cellulose synthesis in cotton fiber, aiming to further validate their predictive power and highlight novel findings. Cotton fiber is composed primarily of cellulose, accounting for over 90% of its composition at maturity (Haigler et al. 2012). Performing a genome-wide analysis of the cellulose synthase (CesA) gene family, we identified 27 CesA genes in the *G. hirsutum* genome and divided them into six classes, consistent with the previous reports (Supplementary Fig. S11) (Zhang et al. 2021c; Wen et al. 2022). Thirteen *GhCesAs* were highly expressed during fiber elongation *via* primary cell wall (PCW) synthesis, and seven were linked to SCW formation after 15 dpa. The remaining seven *GhCesAs* genes exhibit relatively low expression levels throughout fiber development and were considered of unknown function (Figure 4A).

Inspecting the GRN-inferred TF-target relationships involving the fiber development related *GhCesAs*, we next compared how well each GRN method represents these genes and known regulator relationships (Figure 4B). The cGENIE3 network effectively identified all 20 *GhCesAs* as targets and predicted 71 regulatory TFs (Supplementary Fig. S12), resulting in the largest cellulose synthesis subnetwork (Figure 4C). In contrast, cdynGENIE3 identified only 13 *GhCesAs* regulated by 43 TFs, notably missing all of the SCW *GhCesAs* (Figure 4B). Likewise, cCorto identified even fewer (11) *GhCesAs*, again missing all SCW *GhCesAs*, and finding only 8 TFs as regulators (Figure 4B; Supplementary Fig. S13-14). In addition to predicting the greatest number of relationships, cGENIE3 also recovered regulatory relationships verified by prior studies, whereas cdynGENIE3 and cCorto did not. For example, the NAC TFs family genes *GhFSN1_A12* and *GhFSN1_D12* were predicted by cGENIE3 to regulate *GhCesA4* and *GhCesA7*, consistent with their differential expression patterns in *GhFSN1* overexpression lines compared to the wild-type cotton plants that suggest the same regulatory relationship (Zhang et al. 2018). Likewise, *GhWRKY16_D06* was a predicted regulator of *GhCesA7_D7*, aligning with its known role in regulating *GhCesAs* during fiber initiation and elongation (Wang et al. 2021b) (Figure 4C). GO enrichment results showed that the 71 regulatory TFs predicted by cGENIE3 were significantly enriched in plant-type cell wall modification, regulation of secondary cell wall biogenesis, and xylem development (Figure 4F). The results collectively suggested that the cGENIE3 network presents a higher predictive power for cellulose synthesis compared to cdynGENIE3 and cCorto.

Closer examination of the cGENIE3 network revealed two distinct yet interconnected network components (Figure 4C). The smaller component I consists of 11 PCW-related *GhCesAs* (3 *GhCesA1*, 4 *GhCesA3*, and 4 *GhCesA6*) and 27 regulatory TFs. Most of these TFs primarily exhibited peak expression early during PCW synthesis, with 3 exceptions that peaked later. The larger component II includes 7 SCW-related *GhCesAs* (2 *GhCesA8*, 3 *GhCesA7*, and 2 *GhCesA4*) and 38 TFs. Fewer than half of the TFs in this component exhibited concordant expression with their target *GhCesAs.* Among those disconcordant TFs peaking early during PCW formation, the homoeologous pair of top-ranked G2-like TFs described above, *GhMYS1_A10* and *GhMYS1_D10*, were identified (Figure 4C and E). Combining trait association results and expression patterns, *GhMYS1_A10* and *GhMYS1_D10* emerge as potential novel TFs that may positively regulate fiber elongation by promoting PCW formation while inhibiting SCW formation (further explored later; Figure 4C). In addition to more diverse TF expression patterns, component II is enriched for SCW-related genes and is denser and more interconnected than component I, which is enriched for PCW-related genes (Figure 4C-E); this distinction reflects the intricate gene regulatory control underlying the transition from fiber elongation to cell wall thickening. The two components were interconnected through 6 TFs that regulate both PCW-related and SCW-related *GhCesAs*. These findings underscore the utility of GRN interrogation in characterizing key regulators and functions in cotton fiber development.

Regarding At and Dt homoeologous relationships, we identified five TF homoeolog pairs and two *GhCesA* homoeolog pairs in component I, and six TF homoeolog pairs and two *GhCesA* homoeolog pairs in component II. These homoeolog pairs present in the same component accounted for 44.4% (8 of 18) *GhCesAs* and 33.8% (22 of 65) TFs, representing functional conservation or redundant regulatory relationships between homoeologs (Supplementary Table S9). This duplicated nature of allopolyploid gene networks, along with the identification of new master regulators, is discussed next.

### The allopolyploid nature of cotton fiber GRN

Understanding the allopolyploid nature of *G. hirsutum* (2n= 4x = 52; AADD genome) is essential for unraveling the regulatory basis of cotton fiber development. The ascertainment of orthologous-homoeologous relationships among the polyploid A-subgenome (At) and D-subgenome (Dt) genes and their parental A- and D-genome diploids provides a foundation for understanding the evolutionary dimension of duplicated gene regulation during cotton fiber development. We used 22,889 homoeologous pairs that were previously characterized into single-copy orthologous-homoeolog groups (scOGs; each containing a single representative for At and Dt) (Hu et al. 2023) to evaluate the evolutionary outcomes for genes inherited from parental diploids and maintained in duplicate post allopolyploidization. The remaining genes (13,229 At; 15,895 Dt) were categorized into variable-copy orthologous-homoeolog groups (vcOGs), possibly reflecting genetic variation between parental diploids and/or accrued post allopolyploidy. Against this backdrop, we leveraged the network perspectives of gene expression, co-expression, and regulatory interactions, to assess the contributions of the A-versus D-subgenomes for both scOG and vcOG categorization of homoeologous gene pairs during the dynamic process of fiber development.

#### Proportion of fiber-expressed genes

Of the 57,151 fiber-expressed genes (76.3% of the total genome), the A-subgenome contains fewer fiber-expressed genes compared to the D-subgenome (28,004 At vs 29,147 Dt), although this is a higher percentage of the total number of At genes versus Dt (77.53% vs 75.15%; chi-square test *P* = 0.008523). These fiber-expressed genes were further categorized into (1) 19,213 paired scOGs where both At and Dt were expressed; (2) 1,597 unpaired scOGs where only one homoeolog was expressed in fibers (749 At and 848 Dt); and (3) 17,128 vcOG genes (8,042 At and 9,086 Dt) (Table 2: I&II). Gene expressed in fibers represented a significantly higher proportion of the scOG category versus the vcOG category (87.4% vs. 58.5%, respectively; chi-square test *P* = 2.2e-16). Between subgenomes, a higher percentage of vcOG At genes was expressed in fiber versus vcOG Dt genes (60.6% vs 57.2%), while the percentages were comparable for scOGs (87.2% vs 87.7% At and Dt genes, respectively). Thus, the higher percentage of expressed gene content in the A-subgenome was mainly attributable to the higher proportion of vcOGs At genes expressed in fiber.

**Table 2.** Subgenomic contribution to fiber-expressed genes.

|  | **Total** | **At** | **Dt** | **<sup>1</sup>Subgenome contribution** |
| --- | --- | --- | --- | --- |
| **I. All genes in the reference genome** | **74,902** | **36,118** | **38,784** | **At < Dt** |
| scOG genes | 45,778 | 22,889 | 22,889 | - |
| scOG genes | 29,124 | 13,229 | 15,895 | At < Dt |
| **II. Fiber expressed genes** | **57,151** | **28,004** | **29,147** |  |
| **(% of all genes)** | **(76.3%)** | **(77.5%)** | **(75.2%)** | **(At > Dt)** |
| scOG genes: paired | 38,426<br>(83.9%) | 19,213<br>(83.9%) | 19,213<br>(83.9%) | - |
| scOG genes: unpaired | 1,597<br>(3.5%) | 749<br>(3.3%) | 848<br>(3.7%) | At < Dt |
| scOG genes | 17,128<br>(58.5%) | 8,042<br>(60.8%) | 9,086<br>(57.2%) | At < Dt<br>(At > Dt) |
| **III. Genes assigned to co-expression modules** | **25,751** | **12,816** | **12,935** |  |
| **(% of all genes)** | **(34.4%)** | **(35.5%)** | **(33.4%)** | **(At > Dt)** |
| homoeologs in the same module | 12,560; 48.8% | 6,280 | 6,280 | - |
| homoeologs NOT in the same module | 13,191; 51.2% | 6,536 | 6,655 | - |
| homoeologs TF in the same module | 1042; 53.0% | 521 | 521 | - |
| homoeologs TF NOT in the same module | 924; 47.0% | 448 | 476 | - |
| know functional gene in the same module | 52 | 26 | 26 |  |
| know functional gene NOT in the same module | 66 | 33 | 33 |  |
| **IV. Nodes in cGENIE3 network** | **41,757** | **20,578** | **21,179** |  |
| **(% of all genes)** | **(55.7%)** | **(57.0%)** | **(54.6%)** | **(At > Dt)** |
| regulators (TFs) | 1,536 | 773 | 763 | - |
| TFs in scOG pair; % scOG pairs | 1,146; 88.1% | 573 | 573 | - |
| TFs NOT in scOG pair; % scOG pairs | 155; 11.9% | 81 | 74 | - |
| scOG TF | 235 | 119 | 116 |  |
| target genes (TGs): | 41,514 | 20,462<br>(56.7%) | 21,052<br>(54.3%) | (At > Dt) |
| TGs in scOG pair; % scOG pairs | 23,858; 77.9% | 11,929 | 11,929 | - |
| TGs NOT in scOG pair; % scOG pairs | 6,759; 22.1% | 3,291 | 3,468 | - |
| scOG genes; % of all scOG gene | 10,897 | 5,242; 39.6% | 5,655; 35.4% | (At > Dt) |
| **V. Edges in cGENIE3 network** | **149,849** |  |  |  |
| intra-subgenome<br>(average TG number per TF) | 74,701 | At to At:<br>38,704 (50.0) | Dt to Dt:<br>35,997 (47.2) | At > Dt |
| inter-subgenome<br>(average TG number per TF) | 75,148 | At to Dt:<br>40,116 (51.9) | Dt to At:<br>35,032 (45.9) | At to Dt ><br>Dt to At |
| <sup>2</sup> TF: regulatory conservation | 15.8% | 15.7% | 15.8% | - |
| <sup>2</sup> TG: regulatory conservation | 6.4% | 6.3% | 6.5% | - |

<sup>1</sup> Significantly different contribution between subgenomes was shown when Chi-square test $P < 0.05$ .
<sup>2</sup> For given TFs (e.g. At TFs), regulatory conservation measures the percentage of their edges targeting paired At and Dt TGs among all edges. For given TGs, regulatory conservation measures the percentage of their edges regulated by paired At and Dt TFs. Full conservation is represented by 1, while no conservation is represented by 0.

#### Overall expression levels

Comparing expression levels between A- and D-subgenomes revealed a subtle pattern with slightly higher expression of Dt genes (Figure 5A), consistent with previous reports in cotton fibers (Hovav et al. 2008b; You et al. 2023). This expression imbalance was consistently observed for the scOG gene set, whereas the vsOG gene set exhibited the opposite pattern (i.e., higher expression of At genes; Figure 5A). Notably, paired scOGs exhibited the highest expression levels for both At and Dt genes (“scOG pair”: mean TPM of At 18.53 and Dt 19.96), followed by vcOG genes (“vcOG”, At 15.98 and Dt 14.07), and then singleton scOGs with only one homoeolog expressed exhibiting the lowest expression levels (“scOG unpair”: At 1.42 and Dt 1.00).

**Figure 5.**
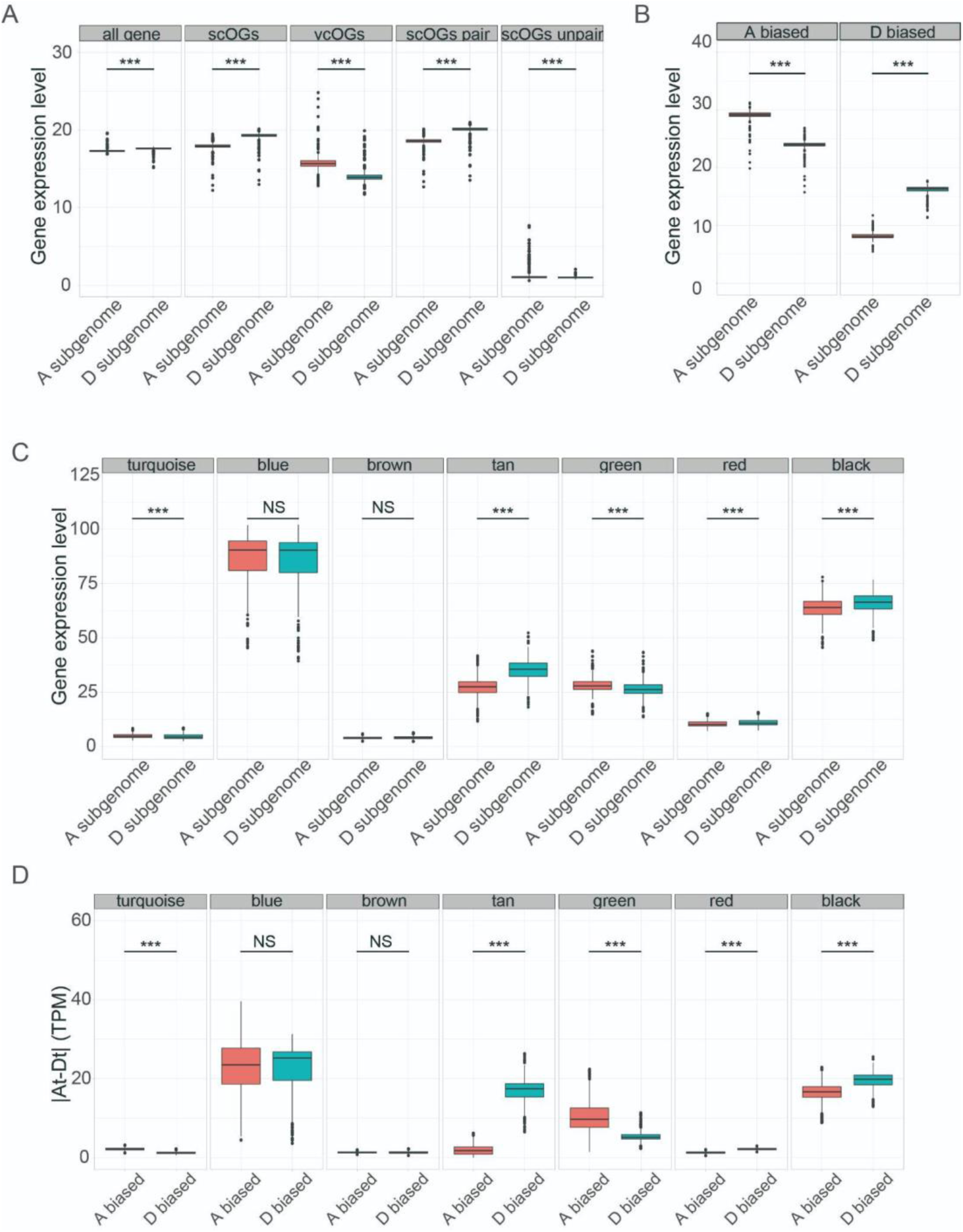
Expression level analysis of homoeologous gene pairs. (**A**) Gene expression levels compared between At and Dt homoeologs for all 57,151 fiber-expressed genes (“all genes”), 22,889 homoeologous pairs characterized into single-copy ortho-homoeolog groups (“scOGs”), the remaining 13,229 At and 15,895 Dt genes uncategorized (“vcOGs”), 19,213 scOGs with both At and Dt expressed in fiber (“scOGs pair”), and 17,028 scOGs with only one homoeolog expressed in fibers (“scOGs unpair”). (**B**) Gene expression levels compared for scOGs pairs exhibiting homoeolog expression bias (HEB). (**C**) Absolute expression differences compared between A-biased and D-biased scOGs. (**C**) Expression comparisons for scOGs present within the same co-expression modules identified by WGCNA. (**D**) Absolute expression differences compared between A-biased and D-biased scOGs in co-expression modules. Statistical significance was determined using a two-sided Wilcoxon rank-sum test. \*\*\**P*< 0.001.

#### Homoeolog expression bias (HEB)

Analysis of HEB, where homoeolog expression statistically varies between duplicates, revealed 8,981 A-biased and 9,153 D-biased pairs among the 19,213 scOG homoeolog pairs, numbers that are not statistically different (*P* = 0.3451), and aligning with previous results (Zhang et al. 2015). Intriguingly, A-biased pairs displayed higher expression levels and larger variation across samples compared to D-biased pairs (Figure 5B). However, D-biased pairs exhibited significantly more expression differences than the A-biased pairs (i.e., Dt-At > At-Dt; Supplementary Fig. S15). This resulted in an overall higher gene expression of scOGs in the D subgenome than the A subgenome, despite the presence of more A-biased versus D-biased pairs.

#### Co-expression modular HEB

The co-expression gene network analysis clustered 25,751 fiber-expressed genes (12,816 At and 12,935 Dt genes) into 20 co-expression modules. Approximately 48.8% of module member genes were paired in modules as homoeologous pairs (6,280 pairs; Table 2: III), indicating substantial functional conservation. The remaining 51.2% of module genes were present in different modules for At and Dt, suggesting functional divergence in terms of co-expression patterns (Supplementary Table S10). Proportions of homoeologous TF pairs in the same module were significantly higher than other homoeologous gene pairs (53.0% vs 48.8%; chi-square test P = 2.2e-16), indicating a higher level of functional conservation between TF homoeologs (Table 2: III; Table S10). Investigating modular HEB for the homoeologous pairs within the same module revealed an absence of significant imbalance of HEB toward either subgenome (Supplementary Table S11). This observation is consistent with the overall pattern of 19,213 scOG pairs. The expression level differences between At and Dt genes across modules (Figure 5C) can be mostly attributed to the expression differences between A-biased and D-biased pairs (Figure 5D; Supplementary Fig. S16). Notably, within the tan module corresponding to fiber elongation, a significantly higher |At-Dt| difference was observed in D-biased then A-biased pairs, implying that the D subgenome might exert a greater effect on fiber elongation than the A subgenome.

#### Subgenomic asymmetry in fiber GRN

Taking the cGENIE3 network as an example, we evaluated the subgenomic contributions to regulatory nodes and edges within the inferred regulatory network. For nodes (Table 2: IV), a higher percentage of At genes was recovered in the network compared to the Dt genes (57.0% vs 54.6%; chi-square test *P* = 0.0005271), and this biased pattern was mainly caused by the target genes (TGs: 56.7% vs. 54.3%; chi-square test *P* = 0.0004837), particularly the scOG ones (39.6% vs. 35.6%; chi-square test *P* = 1.757e-06). The proportion of TFs with both At and Dt homoeologs present in GRN was significantly higher than that of target genes (“TFs in scOG pair” 88.1% vs “TGs in scOG pair” 77.9%; chi-square test P = 2.2e-16). Depending on whether the TF-TG regulatory links were inferred within or between subgenomes, network edges were classified into four categories: two intra-subgenome classes within either subgenome (38,704 At-At and 35,997 Dt-Dt) and two inter-subgenome classes (40,116 At-Dt and 35,032Dt-At) (Table 2: V). The observed ratio of these four edge classes (1.10:1.03:1.15:1.00) significantly deviated from expected proportions assuming full network connectivity from TF to TG nodes (1.01:1.03:1.04:1.00; chi-square test, *P* < 2.2e-16). The intra-subgenome At-At and inter-subgenome At-Dt edges were observed more frequently than expected, indicating a biased regulatory role of At TFs compared to Dt TFs in the fiber gene regulatory network (GRN) (Table 2: V). Finally, we assessed the extent of functional conservation between homoeologs in the GRN, differentiating their roles as TFs or TGs. We observed a significantly higher proportion (15.8%) of edges targeting paired TG homoeologs (i.e., regulatory role as TFs targeting conserved *cis* binding sites) compared to the proportion (6.4%) of edges regulated by paired TF homoeologs (i.e., TGs being regulated by conserved *trans* TF proteins) (Table 2: V; Supplementary Table S12). This suggests that functional divergence between homoeologs in the GRN is more likely to occur in *trans* rather than in *cis*. Consistent patterns were observed in cottonRegNet and other GRNs (Supplementary Table S12).

### Exploring novel regulators of cotton fiber development by GRN inference

We next utilized the best performing cGENIE3 network to identify inter-connections among previously characterized fiber-related genes. Of the curated list of 192 known fiber-related genes (Supplementary Table S3), 154 were present in the network, with 657 directed network edges pointing to them from various TFs. This yielded a seeded network termed kGRN, comprising 432 nodes and 657 edges (Figure 6A, Supplementary Table S13).

**Figure 6.**
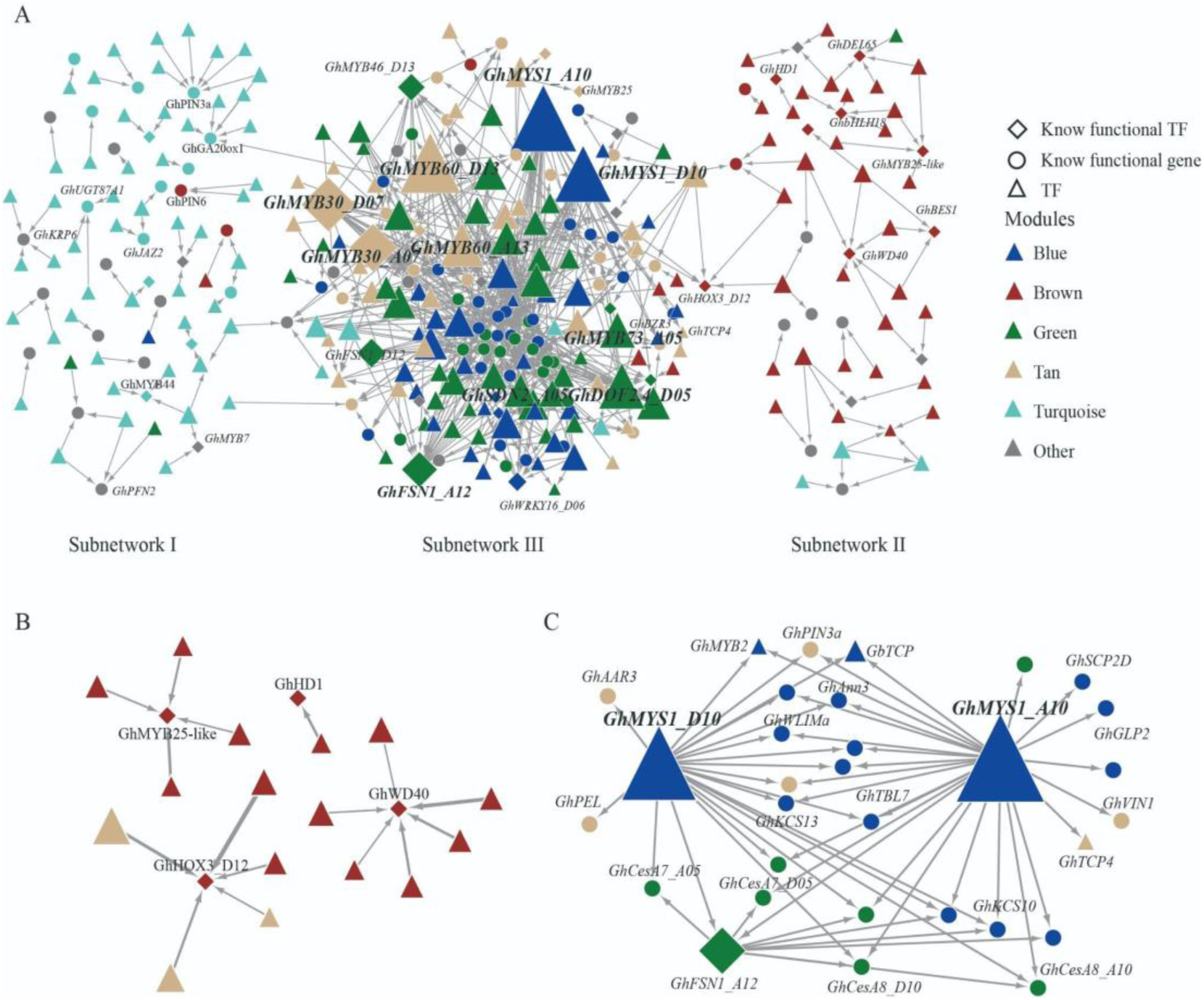
GRN built based on known function genes and their directly regulated TF in fiber. (**A**) GRN of known functional genes and their regulated TFs. Known functional genes and TFs are shown as circles and rhombus, respectively. Different colors indicate the modules where genes and TFs are located in the co-expression network. (**B**) Novel TFs in brown module regulate *GhHOX3*, *GhHD1*, *GhMYB25-like*, and *GhWD40* involved in fiber initiation. (**C**) Network of known functional genes regulated by *GhMYS1_A10* and *GhMYS1_D10*.

Within the kGRN, ten known fiber-related genes function as TFs, regulating other known genes. Eight of these TFs, *GhTCP14*, *GhMYB46_D13*, *GhARF2b*, *GhFSN1_A12*, *GhWRKY16*, *GhGT2*, and a homoeologous pair of *GhMYB30*, have been functionally validated in fiber development (Supplementary Table S14) (Wang et al. 2013, 2021b; Zhang et al. 2018, 2021b; Huang et al. 2019; Tian et al. 2022; Wu et al. 2023). *GhMYB7_A12* and *GhJMJ12_D12*, identified in previous GWAS studies, were significantly associated with fiber strength and/or length (Wang et al. 2017; Liu et al. 2020). Among the remaining 187 TFs with unknown roles in cotton fibers, 97 have *Arabidopsis* orthologs with proven roles in cell wall development or involvement in signaling pathways regulating cotton fiber elongation (Supplementary Table S15), suggesting them as potential candidates for future molecular validation.

By integrating network clustering results with co-expression module annotation, the kGRN was partitioned into three subnetworks (Figure 6A). Subnetwork I, forming a loosely connected periphery on the left, prominently featured co-expressed TFs and target genes from the turquoise module. The target genes of these turquoise module TFs were identified across multiple co-expression modules, suggesting a broad spectrum of regulatory effects amplified by the fluctuating gene expression patterns spanning from 5 to 30 dpa, potentially involving various signaling pathways acting at the inter-modular level (Figure 2A).

Subnetwork II, situated on the right periphery, consists of most of the well-characterized TFs from the brown module, orchestrating key aspects of fiber initiation, including *GhHOX3*, *GhHD1*, *GhMYB25-like*, and *GhWD40* (Figure 6B). Of particular interest is *GhHOX3*, simultaneously regulated by both the brown and tan module TFs (Figure 6B), aligning with its multifaceted role in fiber initiation and elongation functions, respectively (Shan et al. 2014; Qin et al. 2022).

Subnetwork III, centrally located and densely interconnected, weaves together regulatory relationships between TFs and known function target genes in the green, yellow, and blue modules. Among the hub TFs regulating numerous target genes, two, *GhFSN1_A12* (Zhang et al. 2018) and *GhMYB30* (Wu et al. 2023), have been previously characterized. *GhFSN1_A12*, encoding a NAC TF, acts as a positive regulator of fiber SCW thickening by activating a series of known SCW-related genes (Zhang et al. 2018). *GhMYB30,* among the latest characterized members of cotton MYB TFs, was found to regulate cotton fiber development by inhibiting the expression of *GhMYB46*, which was also verified in kGRN (Supplementary Fig. S17) (Wu et al. 2023). Other uncharacterized hub TFs also include *GhMYB73* and a homoeologous pair of *GhMYB60*, offering promising candidates given the well-documented roles of MYB TFs in fiber initiation, elongation, and SCW synthesis.

Focusing on the top hub TFs in kGRN subnetwork III, *GhMYS1_A10* and *GhMYS1_D10* (Figure 6C), represent a homoeologous pair of G2-like TF *MYS1* (*MYB-SHAQKYF 1*). These TFs have known functions in *Arabidopsis* wax biosynthesis and drought tolerance (Liu et al. 2022). Among the 26 targets of *GhMYS1_A10* and 21 targets of *GhMYS1_D10*, 18 target genes were commonly regulated by both, indicating substantial redundancy between homoeologous genes. Among their common targets are several known functional genes including *GhPIN3a*, *GbTCP*, *GhFSN1_A12*, *GhCesA8_D10,* and *GhCesA8_A10* (Figure 6C). In conclusion, known functional genes and their upstream TFs reflect a complex GRN of fiber elongation and SCW synthesis and also helped us identify nine highly connected TFs as candidate regulators of fiber elongation.

### Functional validation of *GhMYS1* reveals its positive role in fiber elongation

Based on the top rankings of *GhMYS1A10* and *GhMYS1D10* in fiber GRNs and their significant trait associations, we selected this homoeologous pair of TFs for functional analysis (Figure 6 and Table S5-6). Comparative expression analysis showed significant upregulation of these genes at 15 dpa in cultivated versus wild cotton and in elite long-fiber versus short-fiber cotton accessions (Figure 7A and B), indicating a potential link with domestication and breeding improvements. Given the 97.29% similarity in the coding regions of *GhMYS1_A10* and *GhMYS1_D10*, VIGS primers were designed to simultaneously silence both genes. VIGS-mediated silencing successfully reduced the expression of both *GhMYS1* genes (Figure 7C), resulting in significantly shorter fibers in pCLCrVA: *GhMYS1* (23.8 mm) compared to pCLCrVA:00 control plants (28.5 mm; *P* =0.001352) (Figure 7D-E).

**Figure 7.**
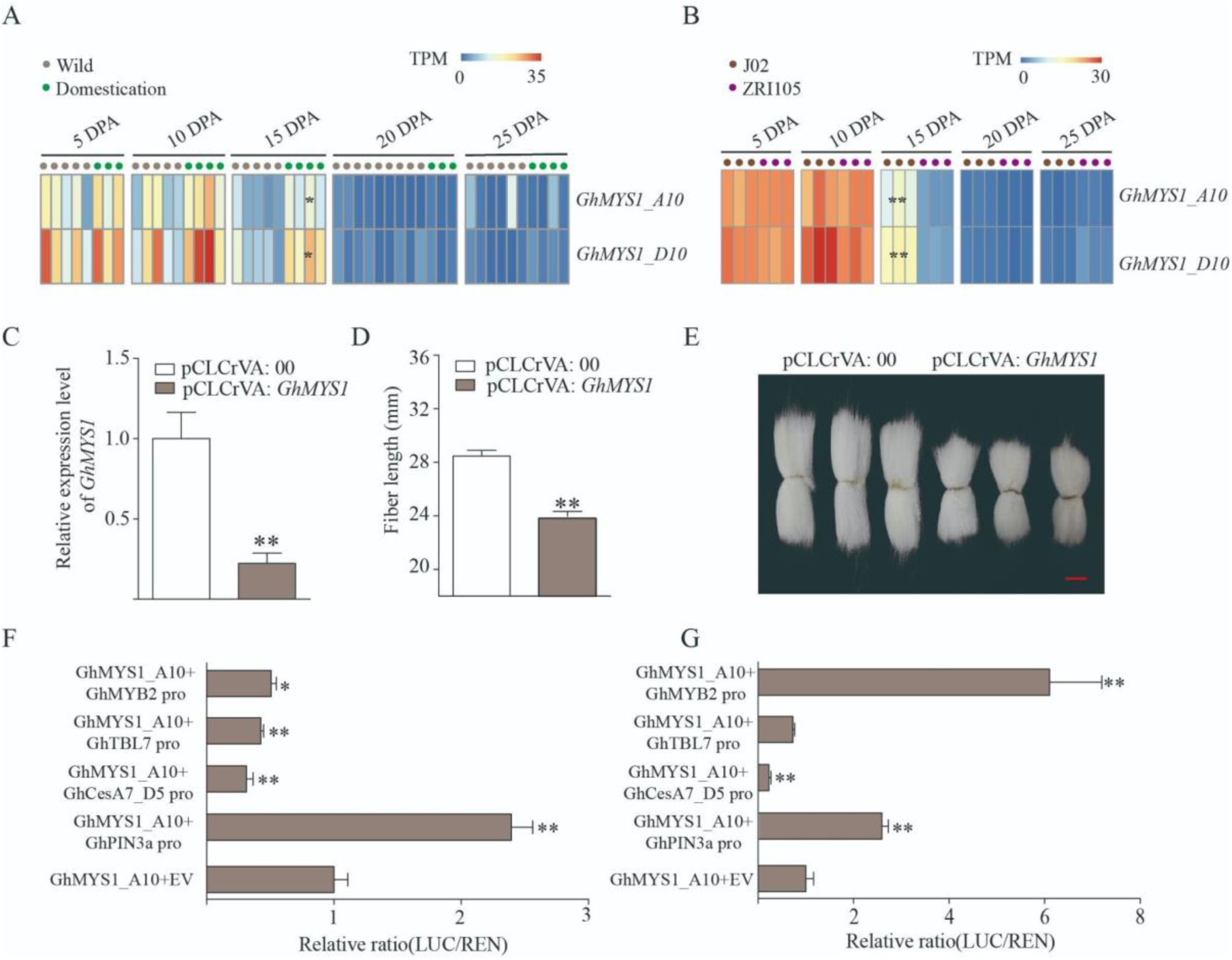
*GhMYS1* positively regulates fiber elongation. (**A**) Expression pattern analysis of *GhMYS1_A10* and *GhMYS1_D10* in wild and domestication cotton accession from 5 to 25 days post-anthesis (dpa). (**B**) Expression pattern analysis of *GhMYS1_A10* and *GhMYS1_D10* in long-fiber (J02) and short-fiber (ZRI105) varieties from 5 to 25 dpa. (**C**) Relative expression levels measured by qRT-PCR showed reduced *GhMYS1* expression in 10 dpa fibers from pCLCrVA: *GhMYS1* cotton plants relative to pCLCrVA: 00 plants. (**D**) Significantly shorter mature fiber length in pCLCrVA: *GhMYS1* versus pCLCrVA: 00 plants. (**E**) Phenotype of mature fibers in pCLCrVA: 00 and pCLCrVA: *GhMYS1* plants. bar = 1 cm. (**F-G**) Transient dual-luciferase (LUC) reporter assay testing interactions between *GhMYS1_A10* (**F**) and *GhMYS1_D10* (**G**), and the promoters of *GhPIN3a*, *GhCesA7_D05*, *GhTBL7*, and *GhMYB2*. Expression of Renilla luciferase (REN) was used as an internal control. Values given are mean ± SD (n = 4). Relative LUC activity obtained with the empty plasmid (none) was set to 1. Statistically significant differences between groups as determined by Student’s t-test. \**P*< 0.05 and \*\**P*< 0.01.

The joint analysis of DAP-seq results and cGENIE3 predictions identified five genes regulated by GhMYS1_A10 and four genes regulated by GhMYS1_D10. Among these, *GhPIN3a* and *GhWLIMa* were common targets regulated by both (Supplementary Table S16). *GhTBL7*, *GhVIN1*, and *GhCesA7_D05* were exclusively targeted by GhMYS1_A10, while *GaMYB2*, and *GbAAR3* was only regulated by GhMYS1_D10 (Supplementary Table S16). Interestingly, *GaMYB2* and three genes including *GhTBL7*, *GhVIN1*, and *GhCesA7_D05* were target genes of GhMYS1_A10 and GhMYS1_D10, respectively, in at least one method of the DAP-seq and cGENIE3. To verify these regulatory relationships, we conducted Dual-luciferase reporter assay (LUC) using the promoter sequences of *GhPIN3*, *GhTBL7*, *GaMYB2*, and *GhCesA7_D05*. The LUC results showed that GhMYS1_A10 could activate *GhPIN3a* while repressing *GhMYB2*, *GhTBL7*, and *GhCesA7_D05*, even though *GhMYB2* was only predicted by GRN (Figure 7F). Consistent with the joint prediction, GhMYS1_D10 activated the expression of *GhPIN3a* and *GhMYB2* (Figure 7G). However, despite GRN and DAP-seq predictions identifing *GhTBL7* and *GhCesA7_D05* as target genes of GhMYS1_D10, respectively, the LUC experiment did not confirm these regulatory relationships, indicating potential false positives (Figure 7G, Supplementary Table S16). The common and discordant regulation of target genes by *GhMYS1_A10* and *GhMYS1_D10* highlights both functional redundancy and differentiation of these homoeologs during fiber development. These findings suggest that *GhMYS1* is a novel transcription factor regulating fiber development, potentially by modulating auxin and positively regulating fiber elongation by suppressing the expression of secondary wall formation-related genes.

## Discussion

### Leveraging GRN inferences for cotton fiber development study

Over the last two decades, conventional molecular genetic analyses have elucidated nearly 200 genes that are important for cotton fiber development, providing valuable insights into the genetic regulation of this process (Huang et al. 2021; Wen et al. 2023). However, previous efforts have often focused on individual genes and limited gene-to-gene inter-connections, resulting in simplified, linear, or limited local networks that fail to capture the complexity of the comprehensive genome-wide GRN governing fiber development. This limitation hinders the full exploration and discovery of the intricate biological networks governing cotton fiber development.

To address this challenge, we leveraged large-scale transcriptome datasets and a wealth of functional gene resources to construct comprehensive genome-wide GRNs for cotton fiber development. We rigorously compared and validated three distinct GRN inference methods against prior knowledge-based regulatory maps, known fiber-related functional genes, DAP-seq data, RNA-seq data from gene perturbation experiments, and phenotypic correlation analyses. This integrative approach demonstrated a carefully crafted, step-by-step process of network evaluation and optimization.

An important consideration in our study was the selection of gene expression thresholds for network construction. Conventional approaches often use generic cutoffs (e.g., greater than 1 RPKM/TPM) (Zhou et al. 2020; Chen et al. 2023), which are prone to exclude transcriptional factors (TFs) and other functional genes with low transcript abundance (Ghaemmaghami et al. 2003; Vaquerizas et al. 2009). In our study, we screened various thresholding options and set the final cutoff at TPM greater than 0 in 30% of samples (see Method or Supplementary Fig. S4), which retained 76.3% of total genes and 72.2% of TFs, ensuring the inclusion of 98.3% of known functional genes crucial for network construction. This approach enabled us to capture a diverse range of genes dynamically expressed across different stages of fiber development, such as *GhHD1* (Walford et al. 2012), *GhMYB25-like* (Walford et al. 2011), *GhWD40* (Tian et al. 2020b), facilitating the identification of the modules related to fiber initiation, even in the absence of fiber samples from 0-4 DPA (Figure 2C, Figure 6B).

GENIE3, an ensemble machine-learning algorithm based on random forests, has demonstrated superior performance in the DREAM4 and DREAM5 GRN reconstruction challenges (Huynh-Thu et al. 2010; Marbach et al. 2012). It has been extensively employed to understand the transcriptional regulation mechanism of plant traits in *Arabidopsis*, rice, wheat, and maize (Walley et al. 2016; Ezer et al. 2017; Huang et al. 2018; Ramírez-González et al. 2018; Shibata et al. 2018; Harrington et al. 2020; Ueda et al. 2020). Given that the fiber transcriptome data in this study consists of 14 time points, dynGENIE3, an adaptation of the original GENIE3 for time series data (Huynh-Thu and Geurts 2018; Balcerowicz et al. 2021), was also used for GRN construction. Additionally, we included another method, Corto (Mercatelli et al. 2020), due to its resemblance to the well-established ARACNe algorithm, which was among the early demonstrated GRN applications known for its ability to infer direct regulatory interactions by eliminating indirect effects (Margolin et al. 2006).

To obtain an approximate “gold standard” for evaluating the performance of GRN predictions, we leveraged the prior knowledge of *Arabidopsis* regulatory interactions (AtRegMap) to assemble a cottonRegMap through orthologous relationships (Wu et al. 2021). Although the limited availability of fiber cistrome data (i.e., TF ChIP-seq or DAP-seq) hinders constructing GRNs directly from empirical evidences of TF-target relationships, we integrated DAP-seq results and transcriptomic data from perturbation experiments for key TFs such as *GhBES1.4* (Liu et al. 2023), *GhWRKY16* (Wang et al. 2021b), *GhMYS1_A10*, and *GhMYS1_D10* to reinforced the reliability of our GRN predictions. In addition to validating the capture of known regulatory relationships, we also considered the inclusion and network centrality of known fiber functional genes, concluding that GENIE3 exhibited the strongest predictive power for known regulatory relationships and key TFs. Notably, these evaluation approaches were applied with appropriate statistical tests (e.g., permutation tests), considering the different network sizes resulting from the three GRN inference methods (with node numbers of 54,237, 25,441, and 56,052 inferred by GENIE3, dynGENIE3, and Corto, respectively).

To exemplify insights gained from this integrative approach, we focussed on the developmentally important process of cellulose synthesis. Although the number of regulator and *GhCesAs* genes captured in networks differed across methods (Table 1 and Figure 5A), cGENIE3 captured the most functionally relevant regulatory relationships in CesA biosynthesis networks. For example, *GhFSN1_A12*’s negative role in suppressing fiber elongation by promoting secondary cell wall (SCW) biosynthesis (Zhang et al. 2018) was evident in its regulation of several *GhCesA* genes involved in SCW formation, including *GhCesA8_D10*, *GhCesA07_D5*, and *GhCesA4_D07*(Figure 4C). Similarly, *GhHOX3_D12*’s involvement in fiber initiation and elongation (Shan et al. 2014; Qin et al. 2022) was supported by its regulation of *GhCesA* genes involved in primary cell wall (PCW) synthesis, such as *GhCesA6_A06* and *GhCesA1_D05-1*(Figure 4C). Notably, homologous genes of SND2 and SND4, key NAC TFs of SCW synthesis in *Arabidopsis* (Taylor-Teeples et al. 2015; Zhong et al. 2021), were also identified as key regulators targeting multiple SCW-related cellulose (Figure 4C). Furthermore, novel regulatory relationships uncovered by cGENIE3, such as GhMYS1_A10’s regulation on *GhCesA7_D5*, were experimentally validated through LUC experiments (Figure 4C, Figure 6C and Figure 7F). These findings underscore the potential of GRN to elucidate molecular mechanisms underlying key TF-gene interactions in fiber development.

### Novel TFs regulating fiber development

GRNs are invaluable tools for predicting the function of TFs. By utilizing the seeded network kGRN of known fiber genes constructed by cGENIE3, we not only confirmed eight previously known TFs (*GhTCP14*, *GhMYB46_D13*, *GhARF2b*, *GhFSN1_A12*, *GhWRKY16*, *GhGT2*, a homoeologous pair of *GhMYB30*) (Wang et al. 2013, 2021b; Zhang et al. 2018, 2021b; Huang et al. 2019; Tian et al. 2022; Wu et al. 2023), but also identified 185 novel TFs regulating known fiber genes (Figure 5A, Supplementary Table S15). Included were a pair of GhMYS1 TFs, GhMYS1_A10, and GhMYS1_D10, that are predicted to regulate 26 and 21 known genes, respectively, and which are ranked highly by prediction scores across all three GRN inference methods, suggesting their important role in fiber development (Supplementary Table S6). This hypothesis is supported by the significant association between fiber traits and gene expression (Supplementary Table S5), where the expression level of this gene pair at 15 dpa fiber is markedly higher in domesticated and elite varieties compared to wild and short-fibered varieties (Figure 7A-B). Experimental validation revealed that silencing *GhMYS1_A10* and *GhMYS1_D10* simultaneously led to a significant reduction in cotton fiber length, underscoring their role in fiber elongation during domestication and breeding processes.

Previous research indicated that MYS1 affects cuticular wax content by down-regulating genes related to wax biosynthesis when overexpressed in *Arabidopsis*, leading to increased contents of primary alcohols, alkanes, and total wax (Liu et al. 2022). With very long-chain fatty acids (VLCFAs) serving as the precursors for wax biosynthesis (Kunst and Samuels 2009) and acting upstream of the ethylene signaling pathway (Huang et al. 2021; Wen et al. 2023), we speculate that MYS1’s role in fiber development involves mediating VLCFA content. Although no significant changes in the content of VLCFAs were detected in the MYS1 overexpression transgenic *Arabidopsis*, MYS1 was co-expressed with several 3-ketoacyl-CoA synthases (KCSs) involved in VLCFA biosynthesis (Liu et al. 2022). Our cGENIE3 results showed that *GhMYS1_A10* and *GhMYS1_D10* simultaneously regulate VLCFA biosynthesis-related genes *GhKCS13* (Shi et al. 2022) and *GhKCS10* (Yang et al. 2023) (Figure 6C), suggesting that *GhMYS1* may affect fiber elongation by regulating VLCFA biosynthesis.

Auxin plays a well-documented positive role in fiber initiation and elongation (Huang et al. 2021; Wen et al. 2023). *GhPIN3a*, an auxin efflux carrier, mediates fiber initiation by establishing hormone gradients in ovule epidermal cells and fibroblast cells (Zhang et al. 2017a; Zeng et al. 2019). Both GRN and DAP-seq results indicated that *GhMYS1_A10* and *GhMYS1_D10* regulate *GhPIN3*, a regulatory relationship further validated by LUC assays (Figure 5C and Figure 7F-G). Additionally, GRN identified four GhCesAs related to secondary wall formation, including *GhCesA7_A05*, *GhCesA7_D05*, *GhCesA8_A10*, and *GhCesA8_D10*, which were regulated by *GhMYS1_A10* and/or *GhMYS1_D10* (Figure 5C). DAP-seq and LUC experiments confirmed the negative regulatory relationship between *GhMYS1A* and *GhCesA7_D05* (Figure 5C and Figure 7E).

Overall, our study validates that *GhMYS1_A10* and *GhMYS1_D10* positively regulate fiber elongation by controlling auxin transport and VLCFA synthesis while inhibiting SCW formation. GRN and DAP-seq results indicate that these TFs regulate numerous genes involved in fiber development (Figure 5C and Supplementary Table S5), suggesting a more complex regulation than previously anticipated. Beyond GhMYS1_A10 and GhMYS1_D10, further exploration of other top-ranking regulators identified by GRN could provide valuable insights for improving fiber quality.

### Asymmetric subgenome contribution to fiber gene expression and network properties

Previous cotton research suggested that the D subgenome exhibits dominant expression (i.e. imbalance of more D-biases than A-biases) and therefore may play a more important role overall than the A subgenome during fiber development and in response to domestication selection (Wang et al. 2017; Ma et al. 2018; Li et al. 2020; You et al. 2023). We note that differences in accessions used, fiber stage, sample numbers and calculation methods among studies have led to varying reports of homoeolog expression bias (HEB), including imbalance favoring D-biased homoeolog pairs (Hovav et al. 2008b; Pei et al. 2022; You et al. 2023) and imbalance (Yoo and Wendel 2014; Zhang et al. 2015; Mei et al. 2021). Against this backdrop of variation, our study found no significant imbalance between A- and D-biased homoeolog expression based on 401 high-quality transcriptome datasets (Supplementary Table S11 and Figure 5A-B).

Beyond the perspective offered by biased homoeolog expression, our analysis explored the nuances of asymmetric duplicated gene expression. Notable findings include a higher number of At than Dt fiber-expressed genes a slightly higher overall transcript abundance of Dt than At genes; and more highly expressed A-biased homoeologous pairs but with lower expression differences (i.e., |At-Dt|) compared to the D-biased homoeologous pairs (Supplementary Table S11 and Figure 5). These subtle and nuanced features and their connections prompted us to speculate that it is the larger expression differences in D-biased homoeologous pairs that contribute to the higher overall transcript abundance of Dt genes, thus leading to the D subgenome exhibiting a disproportionate expression level, which has not been shown in previous studies. These nuanced features enrich our understanding of subgenome contributions to gene expression.

A particularly important methodological consideration is that the analysis of duplicated gene expression, in cotton and other allopolyploid systems (Grover et al. 2012; Bird et al. 2021; Birchler and Yang 2022), typically encompasses single-copy homoeologous gene pairs (or sets, scOGs) derived from the inference of homoeologous relationships. In cotton, even with high-quality genomes and using the latest approaches to orthology inference, such as pSONIC (Conover et al. 2021) and GENESPACE (Lovell et al. 2022), the inferred proportion of single-copy homoeolog groups range from 52% to 73% of the total genomic genic content, meaning that a substantial proportion of genes are missing from analyses of duplicated gene expression patterns. Here we specifically included these variable-copy gene groups (vcOGs) to examine subgenomic contributions and detect previously overlooked patterns. For example, we found that the average expression level of vcOGs in the A subgenome is significantly higher than that in the D subgenome, contrary to the results of scOG (Figure 5A). This finding highlights the importance of considering vcOGs in addition to scOGs when studying gene expression in polyploid systems. It is likely that epigenetic modifications, including DNA methylation and histone modifications, which affect gene expression in polyploid plants (Song and Chen 2015), might also be explored for vcOGs to further our understanding of subgenomic contributions to allopolyploid gene expression. This comprehensive approach will provide a more detailed picture of how gene expression is regulated in polyploid systems.

Perhaps more important than the genic perspective, with respect to the genomic duplication that accompanies allopolyploidy, is that provided by gene co-expression network and regulatory network analyses. These analyses permit the exploration of the joint as well as separate contributions of the A- and D-subgenomes to fiber development, from the standpoint of a more biologically realistic network perspective. Co-expression relationships are often inferred to reflect genes with similar or biologically associated functions (Rhee and Mutwil 2014). Our study shows that scOGs present in the same module account for 48.8% of network genes (Table 2: III), which is higher than the proportions reported for other studies of cotton (Gallagher et al. 2020; Jareczek et al. 2023) and wheat (37.4%) (Ramírez-González et al. 2018). For example, in two previous studies of fiber co-expression gene network construction based on 24 wild and domesticated fiber samples, the proportion of scOGs present in the same module was 20.2-36.1% in *G. hirsutum* and 23.5% in *Gossypium. barbadense* (*G. barbadense*), suggesting that the majority of homoeologous gene pairs are in separate modules in the polyploid network (Gallagher et al. 2020; Jareczek et al. 2023). This discrepancy can likely be attributed to the different RNA-seq samples used. Compared to these earlier studies, our inclusion of more RNA-seq samples, primarily from *G. hirsutum* cultivars, could have resulted in a more connected and denser fiber network due to the effect of domestication, as previously suggested in cotton (Bao et al. 2019; Gallagher et al. 2020) and in other plants (Alonge et al. 2020; Groen et al. 2020). Consequently, we inferred more homoeolog pairs into the same modules, estimating a higher level of functional conservation or closer functional association of homoeologs. Additionally, our larger sample size reduces noise in module assignment, as variable data are more prone to placing homoeologs into different modules. Beyond the overall network structure, our results revealed modular-level features specific to associated functions. For example, the tan and green modules, which were highly expressed during the fiber elongation and SCW thickening stages, showed obvious D and A subgenome biases, respectively. These results further enriched our understanding of the contributions of different subgenomes to fiber development, providing insights that could not be discerned from a single-gene perspective.

Compared to co-expression relationships, TF-TG regulatory relationships inferred by GRNs allow for an examination of subgenomic contributions, including intra-subgenomic interactions (At-At and Dt-Dt) and inter-subgenomic interactions (At-Dt and Dt-At) as previously proposed (Hu and Wendel 2019). This aspect has been explored using three-dimensional genomic interaction (Hi-C) and expression quantitative trait locus (eQTL) methods (Li et al. 2020; Wang et al. 2018). Wang et al. (2018) characterized 3D genome architectures, revealing that inter-subgenomic interactions (At-Dt) accounted for approximately half of all interactions in tetraploid cottons (45.5% in *G. hirsutum* and 47.1% in *G. barbadense*), indicating an equivalent amount of inter- and intra-subgenomic interactions, consistent with our findings. Further, Li et al. (2020) used eQTL analysis on 15 dpa fiber transcriptomes from 251 *G. hirsutum* accessions, identifying 15,330 eQTLs associated with 9,282 genes. They found that the proportion of inter-subgenomic eQTLs was higher in the A subgenome (52.6%) than in the D subgenome (46.5%), suggesting a more prominent regulatory role of At regulators on Dt genes, consistent with our findings. However, they also observed that 44.3% of eGenes in the A-subgenome are regulated by eQTLs in the D-subgenome, whereas only 23.4% of eGenes in the D-subgenome have eQTL regulation in the A-subgenome. This highlights unequal transcriptional regulation patterns between the two subgenomes. An expanded study by You et al. (2023) using fiber transcriptomes from 376 *G. hirsutum* accessions across five time points identified 53,854 cis-eQTLs and 23,811 trans-eQTLs, revealing genetic variants associated with gene expression during fiber development. This larger dataset offers a promising avenue to further delineate inter- and intra-subgenomic regulatory effects and compare them with GRN results. As neither Hi-C nor eQTL analyses directly refined the interaction relationships between TFs and TGs, further analysis integrating eQTL and Hi-C data is needed to obtain TF-TG regulatory relationships and compare them with GRN-based regulatory relationships.

One question of broad interest regarding the functional genomics of allopolyploids is the extent to which duplicated TFs and TGs are functionally conserved in a GRN. A key result emerging from the present work is that the proportion of TG homoeologs simultaneously regulated by any given TF is significantly higher than the proportion of TF homoeologs co-regulating any given downstream genes (e.g., 15.8% vs. 6.4% in cGENIE3; Table 2: V and Supplemental Table S12). This indicates a higher level of conservation in TG promoter *cis*-regulatory sites than in TF *trans* functions. In other words, the *trans*-regulatory roles of TFs diversify faster between homoeologs than does the *cis* landscape of their TG binding sites. This finding is consistent with the experimentally validated notion that *trans*-regulatory mutations have a larger target size compared to *cis*-regulatory mutations in yeast (Siddiq and Wittkopp 2022), hence evolving faster. Further experimental studies in cotton are needed to explore the functional and phenotypic implications of these regulatory variants. For example, in the homoeologous pair of *GhMYS1* genes, DAP-seq results demonstrated both functional conservation and divergence regarding a few target genes with known fiber-related functions. One caveat is that our VIGS experiments can only simultaneously silence both copies due to high sequence identity. Future directions include perturbation experiments targeting individual homoeologs to examine the phenotypic outcomes of disrupting network interactions.

In summary, we constructed comprehensive GRNs using a diverse collection of public RNA-seq datasets for cotton fibers. These rigorously evaluated fiber GRNs enabled us to infer numerous potential regulatory factors controlling fiber development. These include well-studied TFs such as *GhTCP14, GhFSN1_A12, GhWRKY16_D06*, and *GhMYB30*, as well as many TFs with uncharacterized functions. Experimental verification further revealed a key regulatory role of an uncharacterized pair of *GhMYS1* genes in fiber development. Our study reveals subgenomic asymmetries that either accompanied or evolved subsequent to allopolyploidization, including a global expression difference of D-biased homoeolog pairs that underlies the dominant expression of the D subgenome, and further demonstrated multidimensional characteristic of subgenomic asymmetry from the perspective of co-expression and regulatory networks. These findings elucidate the complex gene regulatory network of cotton fiber development, providing insights into the phenomenon of allopolyploidy and offering a resource for exploring genes related to fiber elongation and enhancing cotton fiber quality through breeding.

## Methods

### RNA-Seq data collection and processing

Twelve public cotton fiber RNA-seq datasets comprising 473 samples representing 16 time points of *Gossypium hirsutum* were downloaded from the National Center for Biotechnology Information (NCBI) SRA depository (Supplementary Table S1). Raw reads were preprocessed using fastp (v0.20.1) (Chen et al. 2018) to remove adapters and low-quality reads. Clean reads were aligned to the reference genome *G. hirsutum* var. TM-1 UTX_v2.1(Chen et al. 2020) using Hisat2 (v2.2.1) with default settings (Kim et al. 2015), and transcript abundances were quantified as transcripts per million (TPM) using StringTie (v2.2.1) (Pertea et al. 2015). Dimensionality reduction and visualization of gene expression profiles were conducted through principal component analysis (PCA) and t-distributed stochastic neighbor embedding (t-SNE) in R v4.0.5 (R core Team 2020). The following sample filter criteria were applied to ensure a high-quality dataset: 1) samples were exclusively from fiber tissue, specifically excluding ovular and fibreless mutant samples; 2) samples with a unique mapping rate below 70% were discarded; 3) only uniquely mapped reads were used for TPM calculation; and 4) outlier samples were identified and removed based on PCA and t-SNE.

### Weighted gene co-expression gene network analysis (WGCNA)

A gene co-expression network was constructed using the WGCNA package in R (Langfelder and Horvath 2008) with data from the surviving 401 RNA-seq samples and 57,151 genes. Briefly, the TPM data was used to generate an adjacency matrix based on signed Pearson correlations between all gene pairs powered to an optimized soft thresholding of 28. The adjacency matrix considering gene-to-gene connection strength in isolation was then used to calculate a topological overlap matrix (TOM), which considered each pair of genes in relation to all other genes. Genes with highly similar expression patterns were clustered into co-expression modules, using parameters minModuleSize of 100 and mergeCutHeight of 0.25. Genes belonging to the same co-expression module were assigned the same module color, while genes that cannot be clustered into any of the co-expression modules were labeled grey.

### Construction of gene regulatory networks (GRNs)

Three distinct inference strategies were used to construct fiber gene regulatory networks, including GENIE3 (Huynh-Thu et al. 2010), dynGENIE3 (Huynh-Thu and Geurts 2018), and Corto (Mercatelli et al. 2020). Each method requires both a user-provided list of transcription factors (TFs) and gene expression data to enable inference of directed network connections (edges) from TFs to target genes. A total of 5,048 TFs were identified from the *Gossypium hirsutum* var. TM-1 reference genome (Chen et al. 2020) with PlantTFDB (Jin et al. 2017). Among these, 3,638 fiber-expressed TFs were used as the input TFs to predict targets from all 57,151 fiber-expressed genes. The resulting TF-target predictions were filtered to retain the top one million connections as output GRNs for subsequent analyses, consistent with the thresholding applied in previous studies (Ramírez-González et al. 2018; Harrington et al. 2020). For Corto, which inferred fewer than one million connections, no filtering was applied.

Corto is a correlation-based GRN inference method, implemented as a fast and lightweight R package that resembles the well-established pipeline of ARACNe algorithm (Margolin et al. 2006). Given the normalized TPM data as a gene expression matrix and a list of TFs as centroids, Corto infers direct TF-target relationships through optimized pairwise Pearson correlation. Data Processing Inequality (DPI) on correlation triplets and bootstrapping were applied to evaluate the significance of edges, using the parameters *nbootstraps*=10 and *p*=0.05.

GENIE3 is a machine learning-based approach for GRN inference implemented in R (Huynh-Thu et al. 2010). This method was recognized as the best-performing algorithm in the DREAM4 In Silico Multifactorial challenge (Greenfield et al. 2010) and the DREAM5 Network Inference challenge (Marbach et al. 2012). GENIE3 utilizes the Random Forests tree ensemble algorithm to solve a regression problem for each gene in the given expression dataset, determining how the expression patterns of input TFs predict the expression of the target gene. The importance measure of a TF in predicting the target gene expression serves as the weight for the TF-target regulatory link. GENIE3 was executed using the same gene expression matrix and TF list as input, with default parameters.

Dynamical GENIE3 (dynGENIE3) is an adaptation of the original GENIE3 method that was designed for GRN inference from time series data alone or in conjunction with steady-state data. This semi-parametric model accounts for the dependence between time points by modeling the temporal changes in gene expression with ordinary differential equations (ODEs). In each ODE, the transcription function is learned using a nonparametric Random Forests Model. The fiber gene expression matrix of 57,151 genes and 401 samples was reformatted into two distinct datasets, steady-state and time series, used together as input. The steady-state dataset encompassed 251 samples of 15 days post anthesis (dpa) fibers from Li et al. (2020), focusing on a cultivar population. The time series dataset was constructed using RNA-seq data sourced from other studies (Supplementary Table S1) with at least 3 time points involved: TPM values at each time point were averaged across these studies to obtain the expression profiles spanning 14 time points; genes with a TPM value of 0 in more than two time points were removed, leading to the final inclusion of 1011 TFs and 24,331 other genes. Using both the steady-state and time series data jointly as input, dynGenie3 was executed with default parameters.

### Evaluation of GRN inference

For the performance evaluation of the GRN inference methods, five independent strategies were employed:

I. Homology-based cotton Transcriptional Regulatory Map (cottonRegMap): Serving as a benchmark dataset for validating predicted regulatory links by the above GRN inference methods, this map was constructed by adapting the regulatory prediction approach of PlantRegMap (https://plantregmap.gao-lab.org/) to represent an ensemble list of known regulatory interactions in plants. Briefly, FIMO from the MEME software suite (Bailey et al. 2009) was used to scan TF binding sites in the cotton gene promoters (i.e., 2000 bp upstream of the transcriptional start sites) using a significant threshold of *p*-value <1e-5 with Fisher’s exact test. Regulatory interactions between *Arabidopsis* TFs and cotton gene promoters were assigned if one or more binding sites of a TF were found in the promoter of a gene. Based on the orthologous relationships between 619 *Arabidopsis* TFs and 2,267 *G. hirsutum* TFs (1129 from the At subgenome and 1138 from the Dt subgenome), the TF-target relationships were fully projected onto the *G. hirsutum* genome to form the cottonRegMap, consisting of 53,878,120 TF-target interactions.
II. Cotton TFs with confirmed roles in fiber development: A curated set of 54 TFs with known functions in fiber development was compiled (Supplementary Table 3). Gene set enrichment analysis (GSEA) was used to test if these curated TFs were enriched among the highly ranked TF regulators in each GRN.
III. Physical regulatory relationships based on DAP-seq data: To ground truth the predicted interactions by GRN inference, DNA-affinity purification sequencing (DAP-seq) was performed on a pair of homoeologous G2-like TFs, *GhMYS1_A10* and *GhMYS1_D10*. These TFs were selected based on their consistently high regulator ranking across different GRN inference methods (see Results section for details). Additionally, published DAP-seq data for an *EMS-SUPPRESSOR1v(BES1)/BRASSINAZOLE-RESISTANT1* (*BZR1*) family TF *GhBES1.4* (Liu et al. 2023) was incorporated for validation analysis, which also exhibited high rankings in our GRN inferences. The physical regulatory relationships mapped by DAP-seq were used to validate the GRN prediction by intersecting and significance testing.
IV. RNA-seq analysis of mutants or overexpression lines: We utilized RNA-seq data from TF mutant and overexpression lines to assess the function prediction of candidate TFs in fiber development GRNs. Specifically, RNA-seq datasets for *GhWRKY16* (Wang et al. 2021b) *GhBES1.4* (Liu et al. 2023) reported from previous studies were downloaded. Differential expression analysis was conducted to compare transgenic lines with wild-type controls. The resulting differentially expressed genes (DEGs) were considered potential targets regulated by respective TFs under perturbation conditions, thereby validating the gene targets predicted by GRN. DEGs were identified using DESeq2 (Love et al. 2014) with criteria set at an absolute fold change >1 and the *P*-values <0.05 corrected by the Benjamini-Hochberg method (Benjamini and Hochberg 1995).
V. Fiber traits phenotypic association with gene expression. Corresponding to the 251 RNA-seq samples from 15 dpa fibers (Li et al. 2020), fiber traits from the same natural population were provided by Professor Maojun Wang of Huazhong Agricultural University. The best linear unbiased predictions (BLUPs) of five fiber traits (fiber length, strength, elongation, uniformity, and micronaire value) across the four environments were estimated using the lme4 package in R (Bates et al. 2014). Pearson correlation coefficients were estimated between gene expression levels in 15 dpa fibers and phenotypic variation across the population of 251 cultivars.

### DAP-seq experiments

For the homoeologous TF pair of *GhMYS1_A10* and *GhMYS1_D10*, DAP-seq experiments were performed following the protocol developed by Bartlett et al (Bartlett et al. 2017). Genomic DNA (gDNA) was extracted from 10 dpa fiber of the *G. hirsutum* cultivar TM-1 using the CTAB method. The extracted gDNA was fragmented using a Covaris M220 focused-ultrasonicator (Woburn, Massachusetts, USA) to achieve an average fragment size of 200 bp. These gDNA fragments were used to construct an affinity purification library using the NGS0602-MICH TLX DNA-Seq Kit (Bluescape Hebei Biotech Co., Ltd, Baoding, China). The TF coding sequences were cloned into pFN19K HaloTag T7 SP6 Flexi vector. The TNT SP6 coupled wheat germ extract system (Promega, Wisconsin, USA) was used to express the HALO-tagged TFs in 50 µL reactions, which were incubated for 2 hours at 37 °C. The expressed proteins were directly captured using Magne HaloTag Beads (Promega) and subsequently incubated with the affinity purification library to isolate the TF-DNA binding complexes. The enriched TF-bound gDNA fragments were then eluted from the HaloTag beads, amplified by PCR, and sequenced on the NovaSeq 6000 platform. Two independent biological replicates were conducted for each TF, along with one negative control using a mock DAP-seq library without adding the expressed protein during the HaloTag beads incubation. The DAP-seq raw data have been deposited in the Genome Sequence Archive in National Genomics Data Center, China National Center for Bioinformation / Beijing Institute of Genomics, Chinese Academy of Sciences (GSA: CRA029084 and CRA029060) that are publicly accessible at https://ngdc.cncb.ac.cn/gsa.

### DAP-seq data analysis

Raw DAP-seq reads were pre-processed by removing reads containing adapters and low-quality reads using fastp (v0.20.1) (Chen et al. 2018). Clean reads were aligned to the *G. hirsutum* reference genome using Bowtie 2 (v2.4.5) (Langmead and Salzberg 2012). To identify DAP-seq peaks, MACS2 (v2.2.7.1) peak calling was performed with default parameters (Zhang et al. 2008). Identified peaks from two biological/technical duplicated samples were merged using IDR (v2.0.4.2) to assess the reliability of peak identification (Li et al. 2011). The ChIPseeker R package (v1.40.0) was used for peak annotation in relation to adjacent genes (Yu et al. 2015). Genes with significant peaks (q-value <0.05) within 2000 bp upstream of the transcription start site (TSS) were considered as target genes of the *in vitro* expressed TFs.

### Functional enrichment analysis

Gene functions were annotated based on the eggNOG databases (Huerta-Cepas et al. 2019). Gene Ontology (GO) enrichment analyses were performed using the ClusterProfiler R package (v3.18.1) (Yu et al. 2012). Only GO terms with *P-*values below 0.05 were considered as significantly enriched. GO enrichment results were visualized using aPEAR (v1.0.0) (Kerseviciute and Gordevicius 2023) in R.

### Dual-luciferase (LUC) reporter assay

The 2000 bp promoters of *GhMYB2*, *GhTBL4*, *GhTBL7*, *GhCesA7_D05*, and *GhPIN3a* were cloned using primers listed in Supplementary Table S17 and inserted into the pGreenII 0800-LUC vector. The full-length coding sequences of *GhMYS1_A10* and *GhMYS1_D10* were cloned into the pGreenII 62-SK vector. Resulting plasmids were transduced into *Agrobacterium tumefaciens* strain GV3101, and the LUC reporter assay was performed as previously described (Xie et al. 2017). The pGreen II 0800-LUC and pGreenII 62-SK were used as internal controls. After injecting a mixture of the fusion constructs of pGreenII 62-SK and pGreenII 0800-LUC in a 1:1 ratio into tobacco leaves for 3 days, quantitative analysis of luciferase activity was performed using a Dual-Luciferase Reporter Assay System (E1910, Promega, USA), following the manufacturer’s instructions. All experiments were performed in three independent replicates.

### Virus-induced gene silencing (VIGS) of *GhMYS1*

The cotton leaf crumple virus (CLCrV)-based vectors were used to perform VIGS assays (Gu et al. 2014). To simultaneously silence both *GhMYS1_A10* and *GhMYS1_D10*, a 300 bp coding sequence conserved between homoeologs was designed and inserted into the pCLCrV-A vector to generate the *pCLCrV: GhMYS1* construct. The positive recombinant plasmid of *pCLCrV*: *GhMYS1* and *pCLCrV:00* was subsequently transferred into *Agrobacterium tumefaciens* strain LBA4404 by electroporation. Primers used in vector constructions were listed in Supplementary Table S16. The auxiliary vector pCLCrVB was used to facilitate the intercellular movement of CLCrV DNA. After cultivating *Agrobacterium* colonies containing pCLCrVB, pCLCrV: *GhMYS1*, and pCLCrV: 00 vectors on a shaker at 28 °C for 24 h, *Agrobacterium* cells were collected by centrifugation and resuspended in solution (10 mM MgCl2, 10 mM MES, and 200 mM acetosyringone) to achieve OD600 = 1.2. The *A. tumefacien*s strains containing pCLCrV: *GhMYS1* and pCLCrV: 00 were mixed with pCLCrVB in equal proportions. The resulting mixture was then injected into the cotyledons of 10-day-old seedlings of *G. hirsutum* variety TM-1 using a 1 ml headless syringe. After 24 h of incubation in darkness at 24°C, all plants were transferred to a constant temperature lightroom for cultivation (25°C, 16 hours/day, 8 hours/night). Five plants were injected for each vector, consisting of three biological replicates. The expression of *GhMYS1* was examined in 15 dap fiber of pCLCrV: *GhMYS1* and pCLCrV: 00 cotton plants through RT-qPCR to determine the silencing efficacy.

### Genomic single-copy orthologous-homoeolog groups (scOGs) gene identification

scOGs analysis was carried out by pSONIC software which uses MCScanX and OrthoFinder to infer species pairwise collinearity blocks and identify a high-confidence set of singleton orthologs, respectively (Conover et al. 2021). A total of 22,889 pairs of homologous genes were characterized into scOGs. The remaining 13,229 and 15,895 genes without unique correspondence in At and Dt were named variable copy ortholog groups (vcOGs).

### Subgenomic expression and homoeolog expression bias (HEB) analysis

Because not all of the 45,778 genes placed in scOGs were among the 57,151 fiber-expressed genes, some scOGs were represented in expression data by only the At or Dt homoeolog. Consequently, we further categorized the 22,889 scOGs as either “paired” or “unpaired” based on whether both homoeologs were expressed (scOG paired) or if only one homoeolog was expressed (scOG unpaired). To analyze the expression levels of genes contained within the expressed OGs (vcOGs, scOG paired, and scOG unpaired genes) between the two subgenomes, we compared the average TPM values of 57,151 expressed genes across 401 samples using a two-sided Wilcoxon signed-rank test. For HEB analysis, if the TPM between scOGs in one sample exhibited a more than 2-fold change, the gene pair was identified as a biased homoeologous gene pair in that sample. We utilized a chi-square test and corrected the P value using the Benjamini-Hochberg method to compare the expression bias in 401 samples between the At and Dt subgenomes. When the number of samples with an A or D subgenome bias exceeded the number of samples with a D or A subgenome bias, and FDR ≤ 0.05, we considered that there was an A or D subgenome bias.

## Acknowledgments

We thank members from the Hu Lab and the Ma Lab for helpful discussions. We thank Dr. Vân Anh Huynh-Thu for discussion in GRN construction, the Research IT unit at Iowa State University (https://researchit.las.iastate.edu/) for computational support. This project was supported by the Innovation Program of Chinese Academy of Agricultural Sciences (CAAS-CSIAF-202402) and the National Natural Science Foundation of China (32072111) to GH, the Guangdong Basic and Applied Basic Research (Grant No. 2022A1515110758), and the U.S. Department of Agriculture ARS 58-6066-0-066 NACA “Genomics of Malvaceae” to CEG.

## Author Contributions

G.H. and X.M. conceived the project. X.X. conducted data analysis and wrote the paper, with inputs from JC for ortholog and homoeolog detection. D.Z. conducted data analysis and performed the VIGS and LUC assays. C.E.G., J.F.W., X.M., and G.H., revised the manuscript. All authors read and approved the final manuscript.

## Supplementary data

**Supplementary Figure S1.** Number of RNA-seq samples representing each time point before (left) and after (right) quality control.

**Supplementary Figure S2.** Dimensionality reduction and visualization of gene expression profiles for the 413 public RNA-seq samples passing quality control before removing 12 outlier samples.

**Supplementary Figure S3.** Expression analysis of 192 fiber-related functional genes clustered into three groups

**Supplementary Figure S4.** Criterion testing for filtering fiber-expressed genes.

**Supplementary Figure S5.** Weighted gene co-expression network analysis of 57,151 fiber-expressed genes.

**Supplementary Figure S6.** Enriched GO terms of the seven largest modules as illustrated by an UpSet plot.

**Supplementary Figure S7.** Plant hormone-related GO pathways enriched in the brown (A), tan (B), turquoise (C), and red (D) modules.

**Supplementary Figure S8.** GSEA shows enrichment of known functional TFs in TFs identified by cdynGENIE3(A), and cCorto(B).

**Supplementary Figure S9.** Evaluation of GRN inferences by DAP-seq.

**Supplementary Figure S10.** Evaluation of GRN inferences by RNA-seq.

**Supplementary Figure S11.** Genome-wide characterization of CesA coding genes in *G. hirsutum*.

**Supplementary Figure S12.** Expression pattern analysis of TFs regulating cellulose synthase identified by cGENIE3 in long fiber and short fiber cotton varieties.

**Supplementary Figure S13.** cdynGENIE3 predicted GRN for cotton cellulose synthesis.

**Supplementary Figure S14.** cCorto predicted GRN for cotton cellulose synthesis.

**Supplementary Figure S15.** Absolute expression differences compared between A-biased and D-biased scOGs.

**Supplementary Figure S16.** Gene expression levels compared for scOGs pairs exhibiting homoeolog expression bias (HEB) in co-expression modules.

**Supplementary Figure S17**. GRN of known functional genes regulated by *GhMYB30_A07* and *GhMYB30_D07*.

**Supplementary Table S1.** RNA-seq datasets from 12 studies were used in this study.

**Supplementary Table S2.** Summary statistics of 413 RNA-seq samples passing quality filters.

**Supplementary Table S3.** A curated list of 192 fiber-related genes with known functions.

**Supplementary Table S4.** Significantly enriched GO terms of seven largest co-expression modules identified by WGCNA.

**Supplementary Table S5.** The association analysis between the five fiber traits and the expression in 15 dpa fiber of 77 hub genes identified by cGENIE3.

**Supplementary Table S6.** A comprehensive ranking of TFs based on target gene numbers among cGENIE3, cCorto, and cdynGENIE3.

**Supplementary Table S7.** The association analysis between the five fiber traits and the expression in 15 dpa fiber of 77 hub genes identified by cdynGENIE3.

**Supplementary Table S8.** The association analysis between the five fiber traits and the expression in 15 dpa fiber of 77 hub genes identified by cCorto.

**Supplementary Table S9.** homologous transcription factor and gene pairs in cellulose synthesis-related subnetwork inferred by cGENIE3.

**Supplementary Table S10.** The number of expressed paired and unpaired scOGs in different modules.

**Supplementary Table S11.** Homoeolog expression bias by module.

**Supplementary Table S12.** Estimates of regulatory functional conservation between homoeologs in GRNs.

**Supplementary Table S13.** The information about 432 nodes and 657 edges in kGRN.

**Supplementary Table S14.** The detail of eight known fiber-related TFs in kGRN that directly regulate other known genes.

**Supplementary Table S15.** Functional information of homologous genes in Arabidopsis thaliana of 195 upstream transcription factors in kGRN.

**Supplementary Table S16.** know-function target genes of GhMYS1_A10 and GhMYS1_D10 identified by cGENIE3 and DAP-seq.

**Supplementary Table S17.** The primers used in this study.

